# Functional characterization of putative ecdysone transporters in lepidopteran pests

**DOI:** 10.1101/2022.04.20.488866

**Authors:** George-Rafael Samantsidis, Melina Fotiadou, Savvas Tzavellas, Sven Geibel, Ralf Nauen, Luc Swevers, Shane Denecke, John Vontas

## Abstract

The insect steroid hormone ecdysone plays a critical role in insect development. Several recent studies have shown that ecdysone is transported through **O**rganic **A**nion **T**ransporting **P**olypeptides (OATPs) in insects such as flies and mosquitoes. However, the conservation of this mechanism across other arthropods and the role of this transporter in canonical ecdysone pathways are less well studied. Herein we functionally characterized the putative ecdysone transporter OATP74D from two major agricultural moth pests: *Helicoverpa armigera (cotton bollworm)* and *Spodoptera frugiperda (fall armyworm)*. Phylogenetic analysis of OATP transporters across the superphylum Ecdysozoa revealed that *Oatp74D* is well represented among arthropod species and appeared only at the root of the arthropod lineage. Partial disruption of *Oatp74D* in *S. frugiperda* decreased embryo hatching rate and larval survival, suggesting that this gene is essential for development *in vivo*. Depletion and re-expression of *OatP74D* in the lepidoptera cell line RP-HzGUT-AW1(MG) confirmed the gene’s role in ecdysone import and demonstrated that OATP74D is essential for the transcriptional activation of ecdysone responsive genes including *caspase-3*, implicating this transporter in cell death pathways. Establishment of a simple and robust luciferase assay using the RP-HzGUT- AW1(MG) cell line demonstrated that both HaOATP74D and SfOATP74D are inhibited by rifampicin, a well-known organic anion transporter inhibitor. Overall, this work sheds more light on ecdysone uptake mechanisms across insect species and broadens our knowledge of the physiological roles of OATPs in the transportation of endogenous substrates.

**Author Summary:** The insect steroid hormone ecdysone is critical in regulating many aspects of insects’ life, including development and reproduction. A passive diffusion model was never functionally resolved, but was strongly supported until an organic anion transporting polypeptide was identified to mediate the transport of the hormone. The OATP74D, belonging to the Solute carrier superfamily, has been identified and functionally characterized for the first time in *Drosophila melanogaster*. Although phylogenetic analysis suggests that the *Drosophila Oatp74D* is probably conserved among several insect species, the theory for transporter mediated ecdysone uptake cannot be generalized to all insects without concrete proof. In here we provide functional evidence that the *Oatp74D* of two lepidopteran pest species: *Helicoverpa armigera* and *Spodoptera frugiperda*, is highly required for insect survival and development. Furthermore, we reveal that the OATP74D is necessary to regulate the expression of several ecdysone response genes, including *caspase-3* which is involved in programmed cell death. In addition, we have developed a cell-based platform for screening chemical compounds against the lepidopteran orthologs of *Oat74D* and rifampicin was functionally shown to inhibit ecdysone uptake. Taken all together, our study reveals that *Oatp74D* is conserved among several arthropod species in the ecdysone pathway and given the high necessity for an effective control of these two lepidopteran species, we hypothesized that OATP74D could serve as a possible drug target in those two species.

## Introduction

Steroid hormones are molecules acting as chemical cues governing and coordinating several biological processes in insect physiology, metabolism and development. In hemi- and holometabolous insects the most critical and well-studied steroid hormone is ecdysone [1]. Ecdysone, as a typical steroid hormone, acts either by membrane receptors, which lead to the initiation and modulation of signaling transduction pathways, and/or by their cognate nuclear receptors, which act as transcription factors to selectively regulate target gene expression [2, 3].

During insects’ life cycle, precisely controlled and tightly regulated ecdysone pulses regulate transitions between developmental stages, beginning from the egg hatching stage until pupation [4, 5]. Following secretion from the prothoracic gland, ecdysone is activated to 20E and incorporated into the target cells and initiates a signaling pathway, by binding to the nuclear receptors ecdysone receptor (EcR) and ultraspiracle (USP) which form a heterodimeric transcription factor that initiates a gene expression cascade [1, 6]. Among a large number of tissue specific genes implicated in insect development and regulated by ecdysone, there is a core set of ubiquitously expressed transcription factors induced by the steroid hormone [7]. These are encoded by the early response genes such as the *Eip74A* and *Eip75B*, which in turn lead to the activation of the early late response genes that also express transcription factors like the nuclear receptors HR3 and βFtz-F1, the zinc finger protein Broad and the helix-turn-helix factor E93 [8, 9]. This regulatory hierarchy of genes respond to 20-HE and function as molecular determinants of developmental timing and amplification of the hormone signal in order to ensure successful molting and metamorphosis by initiating diverse and tissue specific-dependent biological processes [10].

In holometabolous insects, a high titer of ecdysone that is released at the final larval stage is necessary for the development of adult structures. While it can promote differentiation and pattern specification through cell cycle regulation in imaginal tissues [11, 12, 13], ecdysone can also initiate programmed cell death in certain larval tissues that will not be required in the adult stage. Secretion of ecdysone commits larvae to pupariation and cessation of growth by orchestrating processes such as proliferation, differentiation, and cell death to ensure the proper development of insects. With regard to cell death, ecdysone regulates the proper activation of programmed cell death (autophagy and apoptosis) in obsolete larval tissues like abdominal muscles, midgut and salivary glands of holometabolous insects like *Drosophila melanogaster* [14, 15], *Bombyx mori* and *Helicoverpa armigera* [16, 17]. Furthermore, ecdysone-induced programmed cell death seems to be necessary for other tissues that undergo remodeling during larval-to-pupal transition, like fat body and certain types of neurons [18]. Transcriptional regulation of genes related with autophagy and apoptosis is governed by ecdysone-induced transcription factors like EcR, BR-C, βFtz-F1, E75A and E75B [18]. Studies in Lepidoptera also indicated that ecdysone is involved in the regulation of both autophagy and apoptosis, which seem to be equally implicated in midgut degradation, and blockage of the pathway causes a severe delay in metamorphosis and lethality [19, 13]. When studying insect cell lines a variety of responses including effects on differentiation and proliferation has been attributed to ecdysone [20].

Although a large part of the regulatory network of ecdysone signaling has been resolved, there was until recently limited knowledge about ecdysone transport mechanisms. For many years, a general theory for simple passive diffusion of steroid hormones prevailed, but this started to be rejected when genetic screens in *Drosophila* identified the presence of transporters that mediate the transport of ecdysone. E23, a member of the ATP-Binding Cassette G (ABCG) protein subfamily, mediates the export of ecdysone in order to regulate the concentration of the hormone into the target cells after executing its function [21]. *Atet*, which also belongs to the ABC protein family, was detected in the prothoracic gland of *Drosophila* and was shown to be involved in importing ecdysone into vesicles which are released by calcium stimulated exocytosis to reach hemolymph [22]. Furthermore, additional work indicated that target cells use an active transport mechanism for ecdysone uptake [23, 24]. Organic anion transporting polypeptide 74d (OATP74D), which belongs to the SLCO family of the Solute Carrier (SLC) transporters, was found to be critical for larval development in *Drosophila*, suggested by the larval arrest observed at the L1 stage when the gene was eliminated, a phenotype that resembled EcR loss of function [24, 25]. Furthermore, OATP74D was found to regulate the ecdysone signaling pathway and to be necessary for ecdysone-dependent gene expression in cultured cells [24]. It is noteworthy to mention that although three additional OATPs mediating ecdysone import have been identified in *Drosophila*, they were dispensable compared to OATP74D (26). Ecdysone was suggested as a substrate for different OATPs, and this has been demonstrated also in mosquitoes which lack an OATP74D ortholog [26]. Considering that human OATPs have been functionally associated with hormone transport [27, 28], it could be suggested that the mechanism of cellular uptake of hormones via OATPs is conserved. However, this requires further functional proof given that a) the OATP transporter family differs significantly among species [29] and b) the case of hormonal transport in insects has been functionally validated in *Drosophila* and more recently in mosquitoes [26], yet limited information exists for other insect species.

*Helicoverpa armigera* and *Spodoptera frugiperda* (Lepidoptera: Noctuidae) are two major agricultural pests damaging several economically important cultivated crops around the world [30, 31]. Most of the control strategies employed to date rely on the use of microbial or small molecule insecticides which are administered orally during the larval stages [31, 32]. Among the several existing compounds, those targeting insect development (such as insect growth regulators, IGRs) are often insect specific . IGRs include the EcR agonists known to activate ecdysone signaling precociously, leading to developmental defects and finally death [32]. Although there are several reports regarding the developmental role of the ecdysone pathway in lepidopteran pests [6], the knowledge about transport and cellular uptake of the steroid hormone in these species is limited.

Here, we tried to analyze the evolution of *Oatp74D* among Ecdysozoa. We further characterized the OATP74D transporters of two lepidopteran species, *H. armigera* and *S. frugiperda*, and showed that the transporter is essential for ecdysone uptake as well as for insect development and survival. Finally, we have developed a cell-based assay, which from a biotechnological aspect could be used for high-throughput screening of inhibitors serving as putative insecticide leads.

## 2. Results

### 2.1 Phylogenetic analysis of OATP74D in arthropod species

Organisms that utilize ecdysteroids, including arthropods and nematodes, taxonomically belong to the group Ecdysozoa and undergo the process of molting during their development. Based on our knowledge that ecdysone transportation is mediated by OATP74D in the arthropod *D. melanogaster* while the nematode *C. elegans* does not carry an *Oatp74D* ortholog [24], we aimed to identify the appearance of this gene during the evolution of Ecdysozoa. For this reason, a rooted phylogenetic tree was constructed using OATPs of representative species of Ecdysozoa (S1 Table). The *Oatp74D* orthologs were clustered into a clade that includes the functionally characterized ortholog of *D. melanogaster* (S1 Table, Fig 1A). The aforementioned clade is well-supported in our phylogenetic analysis, with a bootstrap score of 76 on the divergent branch. This suggests that almost all insects (e.g. *D. melanogaster, Apis mellifera*) along with non-insect arthropods (e.g. mites and crustaceans such as *Tetranychus urticae* and *Daphnia magna*) are represented by one or more copies of *Oatp74D*. A notable exception are mosquitoes which have been previously shown to lack a gene of this clade but contain other ecdysone transporters in this gene family [26]. However, none of the non-arthropod Ecdysozoa (e.g. *Hypsibius dujardini* (Tardigrada)*, Priapulus caudatus* (Priapulida)) was represented in this clade. When considered at the species-level tree (Fig 1B), these data suggest that the evolution of *Oatp74D* appeared somewhere between the divergence of arthropods and priapulid worms such as *Priapulus caudatus*.

**Fig 1.**
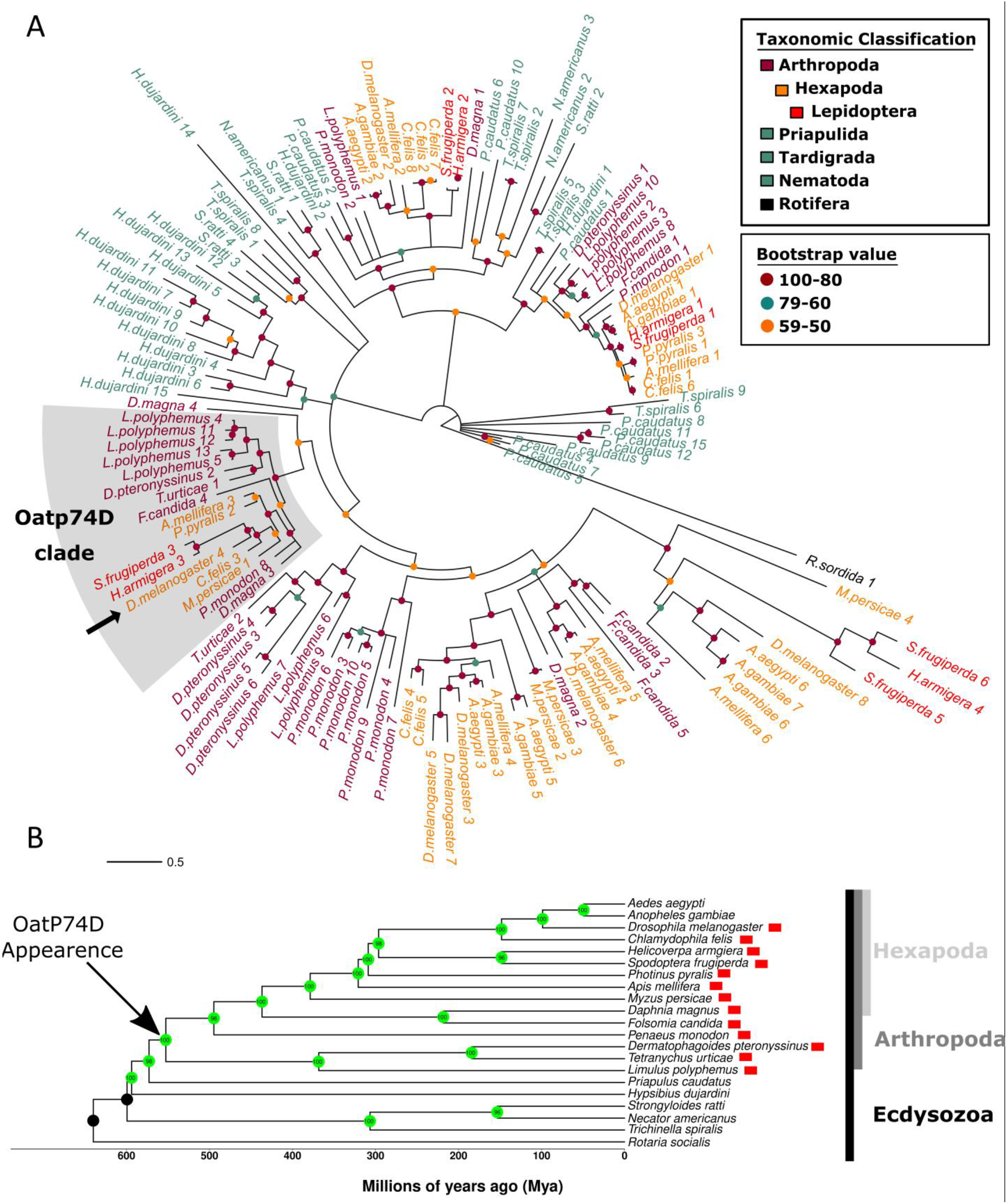
Evolution of Oatp74D in ecdysozoa. (A) Rooted maximum likelihood phylogram constructed using the amino acid sequences of the OATP transporters from ecdysozoan species listed in S1 Table. The tree was rooted using the SLC2 transporter CAF0861090.1 from *R. sordida*. Leaf colors are based on the taxonomic classification of the species, whereas node colors indicate the bootstrap value range. The shaded area indicates the Oatp74D clade including the functionally characterized EcI of *D. melanogaster* (arrow). Scale bar represents an evolutionary distance of 0.5 amino acid substitutions per site. GenBank accession numbers of the proteins used are shown in S1 Table. This tree is depicted with detailed bootstrap values in Figure S6. (B) A species level phylogeny using 1:1 orthologues among species which sampled Ecdysozoa. The x-axis represents estimated divergence times of species in millions of years. Bootstrap support for branches is indicated in green circles at nodes. Greyscale labeling to the right of tip labels indicates clades such as hexapods (light grey), arthropods (dark grey), and ecdysozoa (black). Red boxes indicate the presence of OatP74D in a given species.

### 2.2 *S. frugiperda* OATP74D is essential for larval development

To assess whether OatP74D plays an essential role in lepidopteran development, a CRISPR/Cas9 strategy was employed in order to knock-out the *Oatp74D* of *S. frugiperda* in somatic tisues. Four different sgRNAs were used in order to ensure that a high mutation frequency was displayed at G0 mosaic insects (S1 Fig). As shown in Fig 2A and S4 Table, targeting *SfOatp74D* lead to a significantly lower hatching rate (10%) compared to eggs injected with sgRNAs targeting *SfScarlet* (38%). Furthermore, considering only hatched eggs, a lower proportion of *Oatp74D* injected larvae survived during larval development (25%) compared to the control larvae (86%; Fig 2B). PCR of the targeted region revealed several smaller bands appearing in the *OatP74D* injected larvae while only a single band was obtained from control animals (S1 Fig). This result and subsequent sequencing suggest that deletions of varying sizes were being mediated by the guide RNAs. The positive correlation between mutagenesis of *OatP74D* and lethality is highly suggestive of an essential role for this gene in *S. frugiperda*.

**Fig 2.**
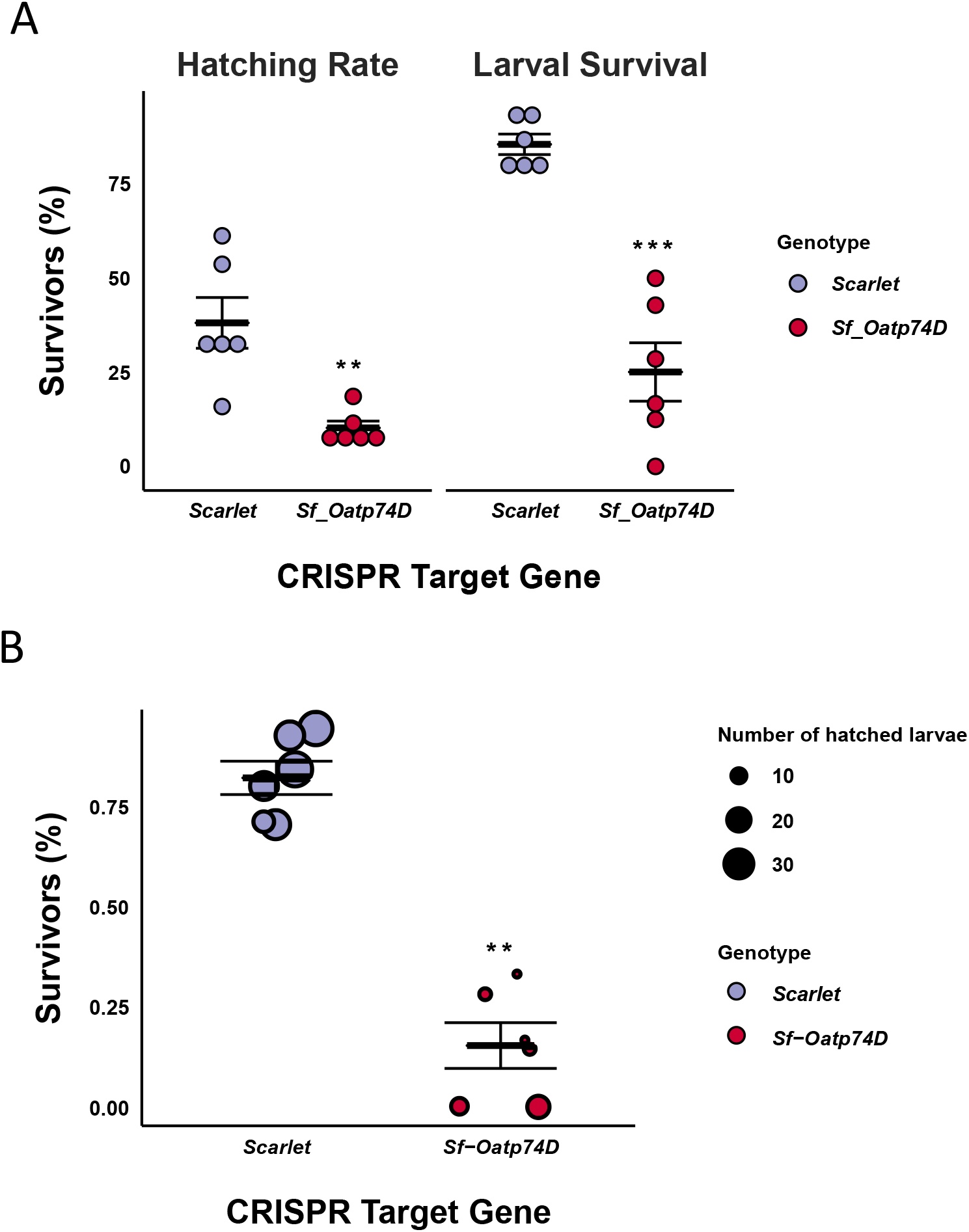
CRISPR mediated disruption of *S. frugiperda Oatp74D.* (A) Lethal stages of S. frugiperda eggs injected with Cas9/sgRNA complexes targeting Oatp74D and Scarlet genes expressed as % of survivors and measured as hatching rate (left panel) and larval survival (right panel). Hatching rate and larval survival are expressed as hatched eggs and larval survivors normalized to total number of injected eggs, respectively. (B) Overall survival rates (as ratio of percentage of survivors against the total number of injected eggs) are illustrated for both Oatp74D and Scarlet genes. The size of each dot is proportional to the number of hathched eggs from each plate. Statistical significance was calculated by using un-paired t-test with Welch’s correction (** p<0.0079 and ***p<0.0003).

### 2.3 *H. zea* OATP74D is necessary for initiation of ecdysone pathway

The high mortality rates at the embryonic stages of *S. frugiperda* made further characterization of the role of lepidopteran OATP74D *in vivo* difficult. Therefore, the HzAW1 cell line was used to analyze the role of lepidopteran OATP74D in the ecdysone pathway. CRISPR-Cas9 was used to target the first exon of *HzOatp74D* and one clonal cell line was generated harboring a 4-bp deletion in the first exon of the gene (S1C Fig). The elimination of these 4bp is predicted to lead to a truncated protein shortly after the start of translation.

#### Knock-out of *HzOatp74D* inhibits differential expression of ecdysone responsive genes

To address if the *HzOatp74D* is implicated in the ecdysone pathway for the regulation of gene expression cascades, four different ecdysone responsive genes were analyzed for their expression following treatment of both wild type and genome-modified HzAW1 cell lines with 1μM of 20-HE for four different time points. Among the four genes analyzed, three of them (*HzEcR*, *HzEip74A* and *HzEip75B*) showed a slight, yet statistically significant, differential expression between the wild type and the knock-out cell lines upon treatment with 1μΜ of 20-HE, at 9hrs and 12hrs of treatment (Fig 3A; S5 Table). *EcR* showed downregulation while *Eip74A* and *Eip75B* were enriched in the KO. Conversely, expression of the *Hr3* gene displayed a more pronounced difference between the HzAW1^WT^ and the HzAW1^ΔOATP74D^ cells. Specifically, treatment of the wild type cells with 20-HE from 6hrs to 24hrs increased the expression of *Hr3* by approximately 9- to 17-folds, with respect to untreated cells, while under the same conditions in HzAW1^ΔOATP74D^ cells the gene was upregulated only by approximately 2-folds at all time points tested. Hence, *HzOatp74D* seems rather essential for the transcriptional regulation of ecdysone responsive genes.

**Fig 3.**
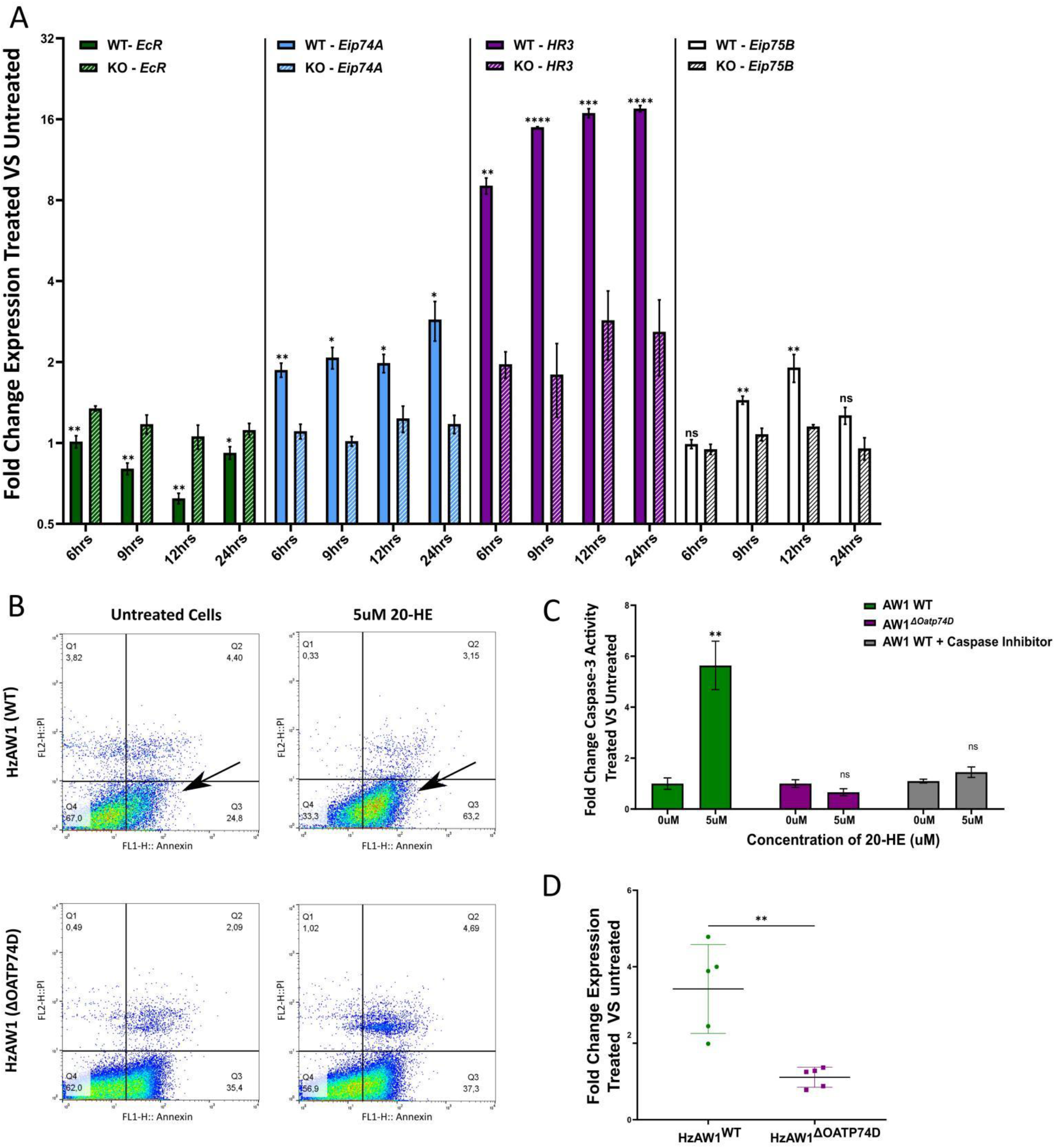
HzOatp74D is necessary for the initiation of ecdysone pathway. (A) Differential expressin analysis of ecdysone responsive genes (HzEcR, HzEip74A, HzEip75B and HzHR3) in HzAW1WT and HzAW1ΔOATP74D cells after treatment with 1μΜ of 20-HE for 6hrs, 9hrs, 12hrs and 24hrs. Fold change expression was calculated as a ratio of the treated versus the untreated cells and expressed as mean of three to six biological replicates. Asterisks indicate statistical significance between the fold change of gene expression between the wild type and knock-out cells for each time point. (B) Flow cytometry analysis of FITC-Annexin V and propidium iodide (PI) stained cells after treatment of both Wild type and monoclonal knock-out cell lines with 5uM of 20-HE for 48hrs; Q1: Annexin-/PI- (live cells), Q2: Annexin+/PI- (early apoptotic cells), Q3: Annexin+/PI+ (late apoptotic cells), Q4: Annexin-/PI+ (dead cells). Flow cytometry plots represents one of the three biological replicates. (C) Fold Change Caspase-3 activity in HzAW1WT and HzAW1ΔOATP74D cells at the same conditions as in B. AW1WT cells were also incubated in the presence of Caspase-3 inhibitor (Ac-DEVD-CHO) during the assay in order to exclude the non-specific cleavage of the synthetic tetrapeptide DEVD. (D) HzCaspase-3 expression analysis in both wild type and knock-out cell lines post treatment with 5uM of 20-HE for 48 hrs. Bars represent the mean ± SE fold change gene expression of the treated versus untreated cells. Asterisks indicate statistically significant differences between the wild type and knock-out cell lines by un-paired t-test, For A: **p=0.0025 and for A:*p<0.0321, **p<0.0046 and ***p<0.0002, ns: non-significant.

### 2.4 HzOatp74D is essential for regulating ecdysone-mediated cell death via caspase-3 activation

One of the physiological functions of the ecdysone pathway at the onset of metamorphosis is the induction of apoptotic cell death [1, 14]. To this end both wild type and knock-out cell lines were treated with 5μM of 20-HE for 48hrs and subsequently assessed for apoptosis using Annexin-PI staining. Treatment of wild type cells with 20-HE for 48hrs increased the percentage of early apoptotic cells (+ Annexin, - PI) by 2.9-folds compared to the untreated wild type cells (Fig 3B). In contrast, the percentage of early apoptotic HzAW1^ΔOATP74D^ cells was at identical levels with the respective negative control (untreated cells harboring the 4-bp deletion at *Oatp74D*).

Apoptotic cell death was further validated via relative quantification of active caspases using a fluorometric assay. As Fig 3C indicates, HzAW1^WT^ exhibited a 5-fold increase of activated caspases upon treatment with 5μM of 20-HE for 48hrs, compared to the untreated wild type cells. However, the HzAW1^ΔOATP74D^ did not display any significant difference when treated with the hormone. Given that the ecdysone pathway regulates the expression of caspases in *D. melanogaster* directly through EcR [14], two different caspases were analyzed for their expression in the treated and untreated wild type and knock-out cells. Relative expression analysis of the two caspases (*caspase-3* and *caspase-8*) of *H. zea* indicated that 20-HE induced the expression of caspase-3 in the wild type cells by almost 3.7-fold compared to the untreated cells (Fig 3D). On the other hand, no difference was observed when HzAW1^ΔOATP74D^ cells were exposed to 20-HE, which further support the Annexin-PI staining and fluorometric assay results. *Caspase-8* was not differentially expressed upon treatment with 20-HE and no difference was observed between wild type and knock-out cell lines.

Taken together these results denote that HzOATP74D is necessary for 20-HE to induce differential expression of ecdysone target genes and apoptotic cell death in the midgut-derived cell line HzAW1.

### 2.5 *H. armigera* and *S. frugiperda* OATP74D are sufficient to rescue ecdysone induced gene transcription in HzAW1ΔOATP74D

To analyze if other lepidopteran orthologs of the ecdysone importer are implicated in 20-HE uptake, a luciferase assay system was implemented similar to that reported previously for the *Drosophila Oatp74D* [24]. Prior to the overexpression of different OATP74D orthologs in cell lines, the assay was first performed with untransfected wild type and knock-out cells to check its robustness. The luciferase assay indicated that HzAW1^WT^ exhibited a significant proportionally increased luciferase activity upon treatment with 0.1μM and 1μM of 20-HE, almost by 3.6- fold and 6.6-fold respectively, compared to the untreated cells (S3B Fig, S4 Fig). On the contrary, treatment of the HzAW1^ΔOATP74D^ with the same concentrations of 20-HE did not induce any increase, keeping the levels of luciferase activity at the baseline (S3B Fig).

Therefore, the HzAW1^ΔOATP74D^ cell line was considered a useful tool for stable expression of other *Oatp74D* orthologs along with the ecdysone responsive firefly luciferase construct, in order to analyze their potency for responsiveness to the steroid hormone. To verify if *HaOatp74D* and *SfOatP74D* expression in the cells, both of them were tagged with a V5 epitope and checked for their expression at a protein level via western blot. As indicated in Fig 4b, lepidopteran OATP74Ds were expressed and identified at the predicted molecular weight, around ∼75KDa. Furthermore, immunostaining of HzAW1^ΔOATP74D^ and Sf9 cells expressing *HaOatp74D-V5* and *SfOat74D-V5* indicated that both proteins are localized at the cellular membrane of the cells as delineated in the bright field overlay (Fig 4a, S3A Fig)

**Fig 4.**
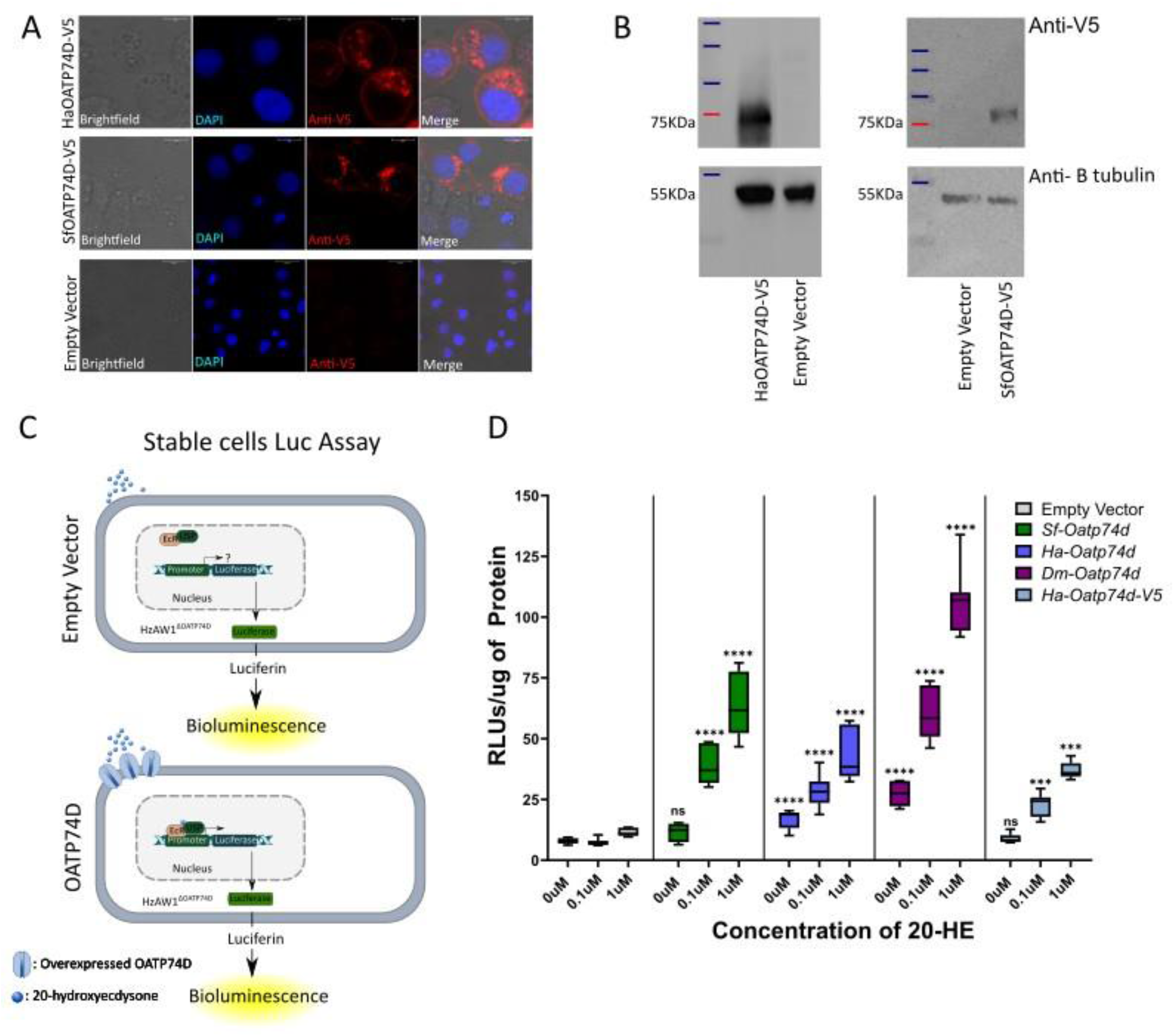
SfOatp74D and HaOatp74D are sufficient to regulate ecdysone induced gene expression in cell cultures. (A) Subcellular localization of SfOATP74D and HaOATP74D tagged with V5 epitope in transiently transfected HzAW1 cells. Blue indicates DAPI that counterstains nuclei, while Red indicates anti-V5, scale bar 20μM. (B) Western blot analysis of SfOATP74D-V5 and HaOATP74D-V5 in HzAW1 stable cell lines. Empty vector stable cells were used as negative control. Top panels represent blots HaOATP74D-V5 (left) and SfOATP74D-V5 (right) along with empty vector stably transfected cells using anti-V5. Bottom panel represent beta-tubulin used as loading control (∼55kDa). (C) Scematic representation of luciferase assay in stable cell lines expressing OATP74D orthologs. (D) Analysis of ecdysone induced luciferase expression in stable cell lines over-expressing SfOatp74D, HaOatp74D, DmOatp74D and HaOatp74D-V5, upon treatment with 0.1 and 1uM of 20-HE for 24hrs. Values are calculated as ratio of Relative luminescence units (RLUs) against the total protein content. Asterisks indicate statistical significant differences between cells overexpressing lepidoptera OATP74D against empty vector, *p=0.0149, ***p=0.0004, ****p<0.0001, calculated with one-way ANOVA followed by post-hoc Dunnett test, ns: non-significant.

Over-expression of *DmOATP74D* was used as a positive control to test if luciferase activity would be induced upon treatment with different concentrations of 20-HE (Fig 4c). A 2.24 and 3.9-fold increase of luciferase activity was observed when cells were treated with 0.1μM and 1μM of the hormone respectively, compared to the untreated cells. Stable cells expressing HaOATP74D displayed a significant increase of luciferase activity by 1.7-fold and 2.54- fold, upon treatment with 0.1μM and 1μM of 20-HE respectively compared to the untreated cells (Fig 4c). Finally, treatment of stably expressing SfOATP74D cells with the same concentrations of 20-HE induced the expression of luciferase by 3.43 and 5.36-fold. Cells transfected with an empty vector did not display any difference upon treatment with 0.1μM and 1μM of 20-HE compared to the untreated cells.

### 2.6 Lepidoptera OATP74D are inhibited by known OATP Inhibitors

Rifampicin and telmisartan, two well-known inhibitors of OATPs, were used in order to test whether OATP74D could be pharmacologically inhibited. Although telmisartan did not impact the function of any of the OATP74Ds (S3C Fig), rifampicin inhibited the ecdysone-induced luciferase activity when tested in stable cells treated with 0.1μM of 20-HE (Fig 5B). In particular, 10μM of rifampicin inhibited SfOATP74D by 30% but did not affect the activity of *Drosophila* and *Helicoverpa* proteins. Conversely, 50μM and 100μM of rifampicin lead to significant reductions of luciferase activity by >50% and >90% respectively, when tested against each of the OATP74D proteins (Fig 5B).

**Fig 5.**
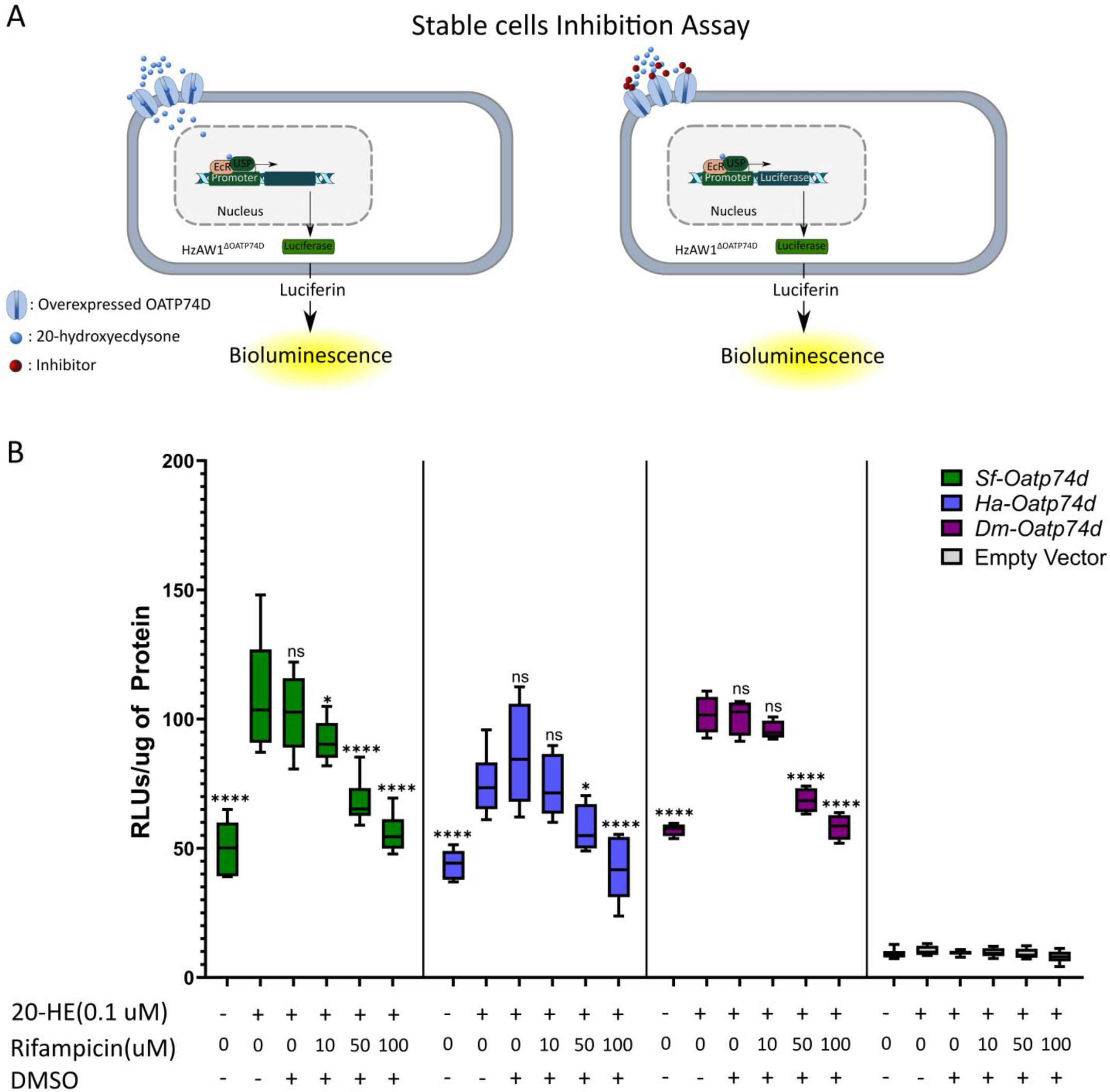
Inhibition analysis of lepidoptera OATP74D by Rifampicin in stable cell lines. (A) Schematic illustration of inhibition assay. (B) Box plot of luciferase assay values when the cell lines were incubated with 0.1uM of 20-HE in the presence of serial concentrations of Rifampicin (10μΜ, 50μΜ and 100μΜ) or DMSO (negative control). Each group is represented by eight different technical replicates. Asterisks indicate statistical significance between the different conditions tested versus cells treated only with 20-HE calculated with one-way ANOVA followed by post-hoc Dunnett’s test, *p<0.048, ****p<0.0001, ns: non-significant.

## 3. Discussion

### 3.1 Phylogenetic analysis of the arthropod OATP protein subfamily

Although the function and phylogeny of OATPs have been extensively studied in mammals, due to their pharmacological significance [27, 34, 51], there is limited information concerning non-mammalian species. Phylogenetic analysis of the OATP family in all orders of Ecdysozoa revealed that *Oatp74D* is well represented among arthropods, while non-arthropod species lack an *Oatp74D* ortholog (Fig 1). Our results are in agreement with previous phylogenetic analysis of this transporter [24] and provide a more detailed view of the presence of *Oatp74D* orthologs in various species of interest, including model species, disease vectors and agricultural pests. Interestingly, apart from identifying *Oatp74D* orthologs in *S. frugiperda* and *H. armigera*, it was found that mosquitoes do not have *Oatp74D* orthologs, although they belong to arthropods (Fig 1). In agreement with this result, a very recent study reports the existence of additional ecdysone importers EcI-2, EcI-3 and EcI-4 in *Aedes aegypti*, with EcI-2 being necessary for development [26]. The fact that orthologs of the latter ecdysone transporters exist in *D. melanogaster*, but have no dominant role in development, exemplifies why further functional characterization of *Oatp74D* was needed to assess its essentiality in the two lepidopteran pests of our interest. Furthermore, the phylogenetic analysis suggested that non-arthropod species appear to have completely divergent clades of *Oatp* transporters which have yet to be characterized (Fig 1).

### 3.2 OATP74D is essential for lepidopteran insect development and survival

Partial disruption of *Oatp74D* in mosaic *S. frugiperda* embryos (Fig 2) had a severe impact on the egg hatching rate. This was not unexpected, since the ecdysone pathway is essential during embryogenesis, as indicated in *Drosophila melanogaster* embryos which seem to express the major biosynthetic enzymes of ecdysone and require EcR-USP nuclear receptors for normal development and survival [52, 53]. Heterologous expression of a dominant negative allele of EcR in a heterozygous mutant background for endogenous *EcR* increased the lethality as well as the penetrance of germ band retraction defects, indicating the necessity of the pathway overall in the development and morphogenesis of embryos [53]. Moreover, null mutant flies of other components of the ecdysone pathway like β*FTZ-F1* and *DHR3* failed to hatch since they exhibited severe defects in ventral nerve cord condensation and an inability to fill their tracheal system with air [9]. Additional studies in lepidopteran species, like *Manduca sexta* and *Bombyx mori*, have documented the expression of ecdysteroidogenic enzymes during embryogenesis [1, 54]. Therefore, the reduced egg hatching rate caused by *SfOatp74D* disruption (Fig 4B) could be explained by its essential role in ecdysone transport. Similar results were observed when disruption of the organic anion transporter EcI-2 of *Aedes aegypti* significantly reduced egg survival [26]. Interestingly, *DmOatp74D* null mutants did not exhibit any significant embryonic lethality, which contradicts with the increased embryonic lethality induced by knocking out *SfOatp74D* (Fig 4B) and *EcI-2* of *Aedes aegypti* [24, 26]. This raises the question whether there is a different mechanism for cellular uptake of ecdysone in the *Drosophila* embryo or the phenotype is masked because of maternal deposition in the eggs of *Oatp74D* mRNA or protein.

High mortality observed in *SfOatP74D* injected individuals at larval stages (Fig 2B) further suggests an essential role for this gene in early larval development. This is in line with *DmOatp74D*, which seems to be essential for the development of larval stages since homozygous mutant flies arrested at L1 stage, failing to molt to the second larval instar [24]. Our results are also consistent with a previous study in *Tribolium castaneum*, in which decreased larval survival and failure of pupation was observed upon silencing of *TcOATP4-C1*, a putative ortholog of the *Drosophila Oatp74D* [55]. Increased larval mortality was also observed in *Aedes aegypti* upon silencing of *EcI-2* which induced 70-80% lethality [26]. It is noteworthy to mention that knock-out of other OATPs of *Drosophila* which were shown to mediate ecdysone uptake *in vitro*, did not impact animal development and survival, indicating the predominant role of *DmOatp74D* [26]. Taken together, *SfOatp74D* is essential for embryo hatching, larval molting and overall survival, although the existence of other lepidopteran OATPs functioning as ecdysone transporters cannot be ruled out.

### 3.3 HzOatp74D is essential for the regulation of the canonical ecdysone pathway and the activation of programmed cell death

In parallel to *in vivo* work, an *in vitro* approach was taken by isolating a clonal cell line that is mutant for *Oatp74D*. Expression analysis suggested that *OatP74D* is necessary for the transcriptional regulation of four different ecdysone-responsive genes, *HzEcR*, *HzEip74A*, *HzEip75B* and *HzHR3*. Differential expression analysis of these genes between the wild type and knock-out cell lines upon treatment with the hormone (Fig 3A) highlighted the role of the *HzOatp74D* gene in the activation of the ecdysone pathway and are in agreement with other studies in which knock- out of *Oatp74D* affected the expression of ecdysone-responsive genes [24, 26].

It is well established that ecdysone is implicated in apoptotic cell death of larval tissues like the midgut and salivary glands during the larval to pupal transition [15, 56]. Previous studies have also indicated the role of certain G-protein coupled receoptors (GPCRs) in the regulation of caspases’ expression as a response to 20-HE induced apoptotic cell death (non-genomic function of ecdysone) [6, 13, 19, 56]. Certain caspases, like the *Drosophila dronc*, *reaper* and *hid* are upregulated at the onset of metamorphosis in tissues like the salivary glands and midgut as a response to the ecdysone pathway through direct binding of the EcR/USP transcriptional complex on the promoter region of these genes [14, 57]. However, the role of *OATP74D* remained unknown in cell death induced by the steroid hormone. Given the lethal phenotype of *S. frugiperda* OATP74D mutants from the early embryo stages, we decided to analyze the involvement of the OATP74D in cell death by using the cell line HzAW1, provided that the knock-out of the gene did not significantly affect the viability of the cells. Treatment of the knock-out and wild type cells with 20-HE indicated a clear difference in the number of early apoptotic cells as well as in the expression levels and activity of *caspase-3* (Fig 3C, 3D), indicating the necessity of the transporter in 20-HE-induced apoptosis. Several studies have documented that the interplay between the genomic and the non-genomic pathway in *H. armigera* is mediated by GPCRs, which modulate gene transcription via regulating EcR and USP phosphorylation [13, 56, 58]. Our results also indicate that OATP74D function is necessary for the activation of the ecdysone pathway and the downstream physiological effects (e.g. triggering of apoptosis), in addition to the possible activation of the non-genomic pathway by 20-HE in the lepidopteran cells.

It must be considered that although the HzAW1 cell line is derived from the midgut tissue of *H. zea* [59], a substantial de-differentiation may have occurred [60], preventing the straightforward transfer of knowledge from *in vitro* to *in vivo*. Nevertheless, the role of OATP74D in the ecdysone pathway could still be resolved, given that regulatory genes of the pathway are expressed in these cells under physiological conditions [46]. A possible explanation to this could be that ecdysone is one of the most important signaling molecules that in very low doses can promote proliferation and growth in insect cultured cells, and therefore most insect cell lines maintain high levels of expression of the ecdysone-related genes [12, 62].

### 3.4 SfOATP74D and HaOATP74D import ecdysone to regulate gene expression

Indirect measurement of ecdysone importation was also accomplished using permanently transformed cell lines that express an ecdysone-responsive luciferase reporter assay together with OatP74D from different insect species. Removal of endogenous *OatP74D* decreased the ecdysone response while re-expression of any ortholog rescued ecdysone import (Fig 4D). To further characterize these transporters, rifampicin and telmisartan were tested for their efficiency to inhibit the function of lepidopteran OATP74D. Both compounds have been previously characterized to act as inhibitors of mammalian OATPs [63, 64, 65]. Although telmisartan had no impact on stable cells expressing OATP74D, rifampicin was shown to inhibit the ecdysone-induced luciferase expression when tested in cells overexpressing SfOATP74D and HaOATP74D, which is indicative that both function as typical OATPs mediating cellular uptake of 20-HE (Fig 5). To the best of our knowledge this is the first time that an ecdysone transporter was shown to be inhibited by a chemical compound, indicating its druggability.

Ecdysone-induced luciferase assays have been used extensively for the characterization of DmOATP74D as well as of other OATPs of both *Drosophila* and *Aedes* in other cell lines such as *Drosophila* S2 and mammalian HEK293 cells [24, 26]. In the case of S2 cells, overexpression of *DmOatp74D* did not exhibit large differences compared to the empty vector in the luciferase assay [24]. Similarly, we found only minor differences when wild type HzAW1 cells were transfected with exogenous *OatP74D* (S4 Fig) suggesting that endogenous *OatP74D* was masking observable measurements [66, 67]. Previous studies performing the assay in mammalian HEK293 cells indicated a clear difference with the negative control given that mammals are void of ecdysone importers [24]. Using HEK293 cells for this assay is more laborious since it requires at least the co-transfection of two major components of the ecdysone pathway, a modified version of the EcR and RXR [24]. It remains unclear which additional components of cell physiology from the relevant organism limit investigations. Thus, HzAW1^ΔΟATP74D^ cells have considerable advantages for the characterization of OATP74D orthologues possessing not only the nuclear receptors necessary for ecdysone response but also better resembling arthropod physiology.

In line with the essentiality of ecdysone transporters and their druggability as shown for rifampicin, it could be hypothesized that n OATP74D could be used as putative insecticide targets. Molting associated endocrine disruption is a known pest control principle, addressed by commercial EcR agonists such as methoxyfenozide which lead to precocious molting [68]. However, targeting OATP74D to block ecdysone signaling could indeed be promising, considering that membrane proteins are possibly more accessible to extracellular compounds compared to cytosolic or nuclear factors. The cell-based screening assay that was developed in this study may facilitate the identification of putative insecticide leads.

## Conclusion

A very old enigma in steroid hormone uptake mechanism has been recently resolved in several case studies in *Drosophila* and mosquitoes. Even when the function of OATP74D in most insect species may be reasonably conserved, the existence of other ecdysone transporters cannot be ruled out even within the same species, as already shown in *Drosophila* [26]. Therefore, unraveling the role of OATPs in other insects’ physiology will further enable understanding of the ecdysone uptake mechanisms. Our study provides useful information about the function of OATP74D in *H. armigera* and *S. frugiperda*, two highly destructive lepidopteran crop pests. The import of 20-HE regulates the initiation of the canonical ecdysone pathway, with *SfOatp74D* being essential for insect survival. This is the first time reported that ecdysone transporters are inhibited by mammalian OATP inhibitors, providing excellent tools for future mechanistic studies. Finally, the HzAW1^ΔOATP74D^ cell line developed in this study can be utilized as a platform for the heterologous expression of other ecdysone transporters for functional studies and screening purposes.

## 4. Materials and Methods

### 4.1 Insects and cell lines

A *Spodoptera frugiperda* population was obtained from Bayer CropScience and was maintained in the lab as a quarantine pest for several generations. The insects were reared at 24±1°C with a 16:8-hour photoperiod on a standard artificial food (based on corn flour).

Two different cell lines were used in this study in order to analyze the role of lepidopteran OATP74D in 20- HE transport. The Sf-9 cell line was obtained from Sigma and maintained as adherent culture in the insect serum free SF900 II SFM (Thermo Fisher Scientific) medium supplemented with 10% heat inactivated fetal bovine serum (FBS, GIBCO, Thermo Fisher Scientific) and 100U/ml of penicillin and 0.1mg/ml streptomycin. The *Helicoverpa zea* midgut cell line RP-HzGUT-AW1(MG) (referred hereafter as HzAW1) was a generous gift by Dr. Cynthia L. Goodman (Biological Control of Insects Research, U.S, Department of Agriculture, Agriculture Research Service). The cell line was routinely maintained as adherent culture in Excell 420 insect serum-free medium (Sigma Aldrich), supplemented with 10% heat-inactivated fetal bovine serum (FBS, GIBCO, Thermo Fisher Scientific) and 100U/ml of penicillin and 0.1mg/ml streptomycin. Both cell lines were kept in a humidified incubator at 27°C.

### 4.2 Phylogenetic analysis of OATP74D in arthropod species

#### Construction of the OATP gene family tree

The evolutionary history of *Oatp74D-*like genes was characterized by phylogenetic analysis using a representative subset of species from the Ecdysozoa taxonomic group (S1 Table). The reference gene annotations and proteomes of these species were downloaded from the National Center for Biotechnology Information (NCBI) and filtered in order to contain only the longest amino acid isoform per gene. Then, the *SLC_id* pipeline [38] was applied on the filtered proteomes to select the OATP (SLCO aka SLC21) transporters of these species. Multiple sequence alignment was performed for the amino acid sequences of the identified OATPs and the outgroup *R. sordida* CAF0861090 SLC2 transporter (S1 Table) using Mafft v7.450 [33] under the default parameters. The outgroup was selected based on the classification of the Major Facilitator Superfamily (MFS) [34]. The produced alignments were automatically trimmed using TrimAl v1.4.rev22 -automated1- heuristic method [35]. Finally, the phylogenetic tree was built under the maximum likelihood optimality criterion by making use of RAxML-NG v. 0.9.0, with the parameters --bs-trees autoMRE{500} for 500 bootstraps and --model LG+G8+F for model specification [36]. The tree was visualized using the Ape package in R [37]. The orthogroup of *Oatp74D* was identified based on the functionally characterized orthologous gene from *D. melanogaster* [24].

Two paralogs of *Oatp74D* from *S. frugiperda,* LOC118271297 and LOC118278121, were identified in the Oatp74D clade. The aforementioned genes were manually inspected to assess if a gene duplication has taken place in the species. The assembly of *S. frugiperda* used in this analysis, GCF_011064685.1, was compared to two different *S. frugiperda* assemblies, available in the NCBI under the accession numbers GCA_012979215.2 and GCA_019297735.1 and the latter was found to include only one copy of the gene. Thus, from the two *Oatp74D* genes identified in the phylogenetic analysis, only 118271297 was retained for the downstream analysis, having higher percentage of identity to the corresponding *Oatp74D* genes of the two other assemblies. The tree was reconstructed as described above, after excluding LOC118278121 from the multiple sequence alignment.

The gene level phylogeny was complemented by a species level phylogeny using representative species from Ecdysozoa (S1 Table) as performed previously [38]. Briefly, one-to-one orthologs were obtained using Orthofinder [39] with the filtered proteome files as inputs with default parameters. All orthologs from each orthogroup were aligned using Mafft and trimAl as in the gene level phylogeny [35, 40]. These alignments were then concatenated into a single alignment which was used as an input for maximum likelihood tree building with RAxML-NG v. 0.9.0 -- bs-trees autoMRE{200} for 200 bootstraps and --model LG+G8+F for model specification. Visualization was accomplished using the ggtree package in R [41].

### 4.3 Construct preparation

#### Plasmids for transient OATP74D over-expression in insect cell lines

The open reading frames of *SfOatp74D* (Gene ID: 118271297, 2109bp) and *DmOatp74D* (Gene ID: 39954, 2460bp) were PCR amplified using Phusion polymerase (NEB) from cDNA templates of 3^rd^ instar larvae of *S. frugiperda* and adults of *D. melanogaster* respectively. The primer pairs used for PCR amplification were Sf-OATP74D-XbaI-F/Sf- OATP74D-NotI-R and Dm-OATP74D-XbaI-F/Dm-OATP74D-NotI-R (S2 Table), respectively. The PCR reactions for both genes were performed as follows: 98°C for 30sec initial denaturation, followed by 30cycles of 98°C for 10sec, 63°C for 30sec, 72°C for 1min10sec, followed by final extension at 72°C for 5min. Both PCR products were purified with a PCR clean-up kit (Macherey-Nagel) according to manufacturer’s instructions. Both fragments were cloned into the shuttle vector pGEM-T easy (Promega) and verified by Sanger sequencing. The *HaOatp74D* (2136bp) ORF was synthesized *de novo* (Genscript, Piscataway, NJ) based on the alignment of both NCBI reference sequence and the *de novo* transcriptome assembly of *H. armigera* [42]. The newly synthesized sequence was subcloned between the BamHI and NotI restriction sites of pFastBac1 vector. The *SfOatp74D* and *DmOatp74D* were finally cloned in between the XbaI and NotI sites of the lepidoptera specific expression vector pBmAc3 [43] while *HaOatp74D* was cloned between BamHI and NotI sites.

#### Plasmids for stable cell line generation

The pEIA vector [49] was modified with the Gibson assembly methodology in order to replace the BmNPV-IE1 ORF with Puromycin N-acetyltransferase (PAC). The primers used to amplify the pEIA plasmid were pEIA-Fgibson and pEIA-Rgibson (S2 Table) and the PCR reaction was performed using Phusion polymerase (NEB). The ORF of the puromycin resistance gene was amplified using Phusion polymerase and pEA-PAC as a template and the primer pair used for the PCR reaction were PAC-Fgibson-AscI/PAC-Rgibson-NcoI (S2 Table). Both primers introduce the restriction sites of the unicutters AscI and NcoI to facilitate cloning of any other gene of interest downstream of the BmNPV-iE1 promoter. Both PCR products were used for constructing the final vector with Gibson assembly Master Mix (NEB), according to the instructions of the manufacturer. The final vector was verified by sequencing (Genwiz, Germany) and named as piE1:puro-BmAc3. To replace puromycin N-acetyltransferase with the Zeocin resistance gene (*Sh ble*), the pPICZa vector was digested with NcoI and EcoRV. The generated 439bp fragment was cloned into the vector piE1:puro-BmAc3 digested with AscI, followed by treatment with Klenow fragment (Minotech) and subsequent digestion with NcoI. The ORF of *SfOatp74D*, *DmOatp74D* and *HaOatp74D* were cloned into the final vector (piE1:Zeocin-BmAc3) using the same strategy as used in the case of pBmAC3 vector.

To tag both *SfOatp74D* and *HaOatp74D* with a V5 epitope (GKPIPNPLLGLDST) at the C-terminus of the protein, both ORFs were amplified with PCR using the primer pairs Sf-OATP74D-XbaI-F/Sf-OATP74D-BspEI-V5-NotI-R and Ha- OATP74D-BamHI-F/HaOATP74D-BspEI-R, respectively. The SfOATP74D insert was cloned in between XbaI and NotI sites of piE1:Zeocin-BmAc3 vector, harboring a BspEI restriction site upstream of the V5 epitope sequence. The *HaOatp74D*-*V5* PCR fragment was cloned between the BamHI and BspEI of the pBmAc3-SfOATP74D-V5 vector. A linker sequence (Gly-Ser-Gly) was used to separate the C-terminus of each of the two proteins with the V5 epitope. All plasmids generated in this study can be found in S3 Table.

### 4.4 CRISPR mediated Knock-out of Oatp74D in *S. frugiperda*

In order to somatically disrupt the *Oatp74D* gene *in vivo*, CRISPR-Cas9 was performed by injecting *S. frugiperda* eggs according to a previously established protocol [44]. Briefly, egg batches were collected shortly after the onset of the scotoperiod and transferred to double sided tape using a whetted paintbrush [26]. Eggs were then injected under air-dry conditions with a solution containing 300ng/μl of recombinant Cas9 Nuclease (NEB) and 100ng/ul each of four sgRNAs targeting the first exon of the *Oatp74D* gene (S2 Table). Two days post-injection, wheat powder was sprinkled on top of the tape, which prevented the larvae from sticking once emerged. Survivorship and the number of days until pupation were measured across the lifespan of the emerging larvae. DNA samples obtained from healthy and weak larvae were sent for amplicon sequencing (GeneWiz) using primers flanking the four sgRNA cut sites (S2 Table). As a control for normalizing lethality due to technical handling during microinjections, *S. frugiperda* eggs were injected with sgRNAs targeting the *Scarlet* gene, which does not impact insect development and survival [45].

### 4.5 CRISPR mediated knock-out of HzOatp74D in HzAW1 cell line

A CRISPR-Cas9 strategy was employed to knock-out *HzOatp74D* in the HzAW1 cell line. Several CRISPR targets were identified in the first exon of the gene based on the *de novo* transcriptome assembly of HzAW1 cell line [46], using the online version of the target finder chopchop [47]. Two different target sequences were selected displaying the minimal predicted off-target effects and the highest predicted efficiency. Single guide RNA sequences were annealed as single stranded oligos (S2 Table) and ligated into the CRISPR vector pBmAc3:Cac9-HaU6:1 [43] following digestion with BbsI.

The HzAW1 cell line was co-transfected with the two sgRNA expressing vectors and the pEA-PAC plasmid at a molecular ratio of 10:10:1. Specifically, one million cells were seeded in 6-well plates and co-transfected with 1μg of total DNA using the ESCORT IV transfection reagent (Sigma) following the instructions of the manufacturer. To positively select the transfected and possibly mutant cells, selection with 25ug/ml of puromycin was carried out for 10 days. Genotyping of the two generated cell lines was performed with PCR using primers flanking the targeted region (HzOATP74D-F-5UTR and HzOATP74D-R-exon1, S2 Table) yielding a fragment of 912bp corresponding to the wild type allele. PCR reactions were performed using Taq DNA polymerase (EnzyQuest, Greece) on genomic DNA extracted from both transfected cell lines with DNAzol reagent (Molecular Research Center); the conditions of the PCR were as follows: 95°C for 3min initial denaturation, followed by 30cycles of 95°C for 30sec, 50°C for 30sec, 72°C for 30sec, followed by final extension at 72°C for 5min. The combination of sgRNAs yielded two distinct products corresponding to the wild type (912bp) and the mutated allele (∼800bp) (S2B Fig). Each of the generated PCR fragments was purified and sequenced to validate the existence of mutated OATP74D isoforms.

Once the existence of mutated OATP74D alleles were verified by Sanger sequencing, single cell cloning was initiated to isolate a monoclonal line encompassing a unique isoform of mutated OATP74D gene. Limiting dilution method was employed in order to isolate clonal cell lines in a 96 well plate. From the 96 well plate, 16 wells were found to contain colonies of cells proliferating and only one of them were found to bear a single mutated OATP74D isoform. The monoclonal cell line was subsequently scaled up and used for downstream assays.

### 4.6 Expression analysis of Ecdysone responsive genes in HzAW1WT and HzAW1ΔOATP74D

The significance of OATP74D in ecdysone signaling of lepidoptera species was examined by analyzing the expression of four different ecdysone early response genes (NCBI GeneIDs: 110373773 (*HaHr3*), 110369974 (*HaEcR*), 110374646 (*HaEip74A*), 110370041 (*HaEip75B*)) in the parental HzAW1 cell line bearing the wild type allele of *Oatp74D* and the monoclonal cell line bearing the mutated isoform of *Oatp74D* (referred hereafter as HzAW1^WT^ and HzAW1^ΔOatp74D^ respectively). Wild type and *HzOatp74D* knock-out cell lines were treated with either 1 μM of 20-HE or the solvent (0.01% Ethanol) for four different time points (6, 9, 12 and 24 hrs). RNA was extracted from each group using Trizol reagent (MRC), according to the instructions of the manufacturer, combined with DNAse treatment using the Turbo DNA free kit (Invitrogen). One μg of total RNA was used for first-strand complementary (cDNA) synthesis using specific primers with the Minotech RT kit (Minotech). qRT-PCR was performed on a CFX connect real-time PCR detection system (Bio-Rad) using the KAPA SYBR fast qPCR Master Mix kit (Kapa biosystems). The reactions were carried out using the following conditions: 95°C for 3min, followed by 40 cycles of 95°C for 10sec and 60°C for 45sec. All primers (S2 Table) were designed based on the *de novo* transcriptome assembly of the HzAW1 cell line [46]. The efficiency of PCR for each primer pair was assessed in 5-fold dilution series of pooled cDNA samples. The experiment was performed using three biological replicates and two technical replicates. Fold change expression was calculated as previously described [48]. Relative expression was normalized against the housekeeping genes *HzGadph* and *HzRps3a*. All primers that are used for the qRT-PCR experiments are summarized in the S2 Table. The fold change of the relative expression data of qRT-PCR between the wild type and *Oatp74D* knock-out cell lines were analyzed for significance using un-paired t-test.

### 4.7 Analysis of ecdysone induced cell death in HzAW1 cell lines

In order to assess the role of OATP74D in ecdysone-mediated cell death, 4x10^5^ cells of both HzAW1^WT^ and HzAW1^ΔOATP74D^ cells were seeded in 6 well plates and treated with 5μM of 20-HE for 48hrs. Each condition consisted of three biological replicates. The cells were harvested and 10^5^ cells from each replicate were used for Fluorescence Activated Cell Sorting following staining with Annexin-PI (BD Pharmigen). The rest of the cells were used for protein and RNA extraction. Caspase-3 activity was calculated using the Caspase-3 assay kit (BD biosciences) following the instructions of the manufacturer. Furthermore, the Ac-DEVD-CHO (BD Biosciences) was used as a potent Caspase-3 inhibitor to validate that fluorescence is mediated by caspase specifically and not by other serine proteases like cathepsins. Fluorescence was measured using the spectramax plate-reader with an excitation wavelegnth of 380nm and an emission wavelength range of 420-460nm (with 5nm increment). For each condition three biological and two technical replicates were used.

Extracted RNA from each sample was used for cDNA synthesis for qRT-PCR to analyze the expression of the genes *HzCaspase-3* and *HzCaspase-8* as previously described (NCBI Gene IDs: 110374006 (*HaCaspase-3*) and 110369675 (*HaCaspase-8*)). Primer sequences used for both genes are shown in the S2 Table. Relative expression was normalized against *HzGadph* and *HzRps3a*.

Calculation of the proportion of apoptotic cells after treatment with 20-HE with Annexin-PI staining was conducted using FlowJo V10 software (BD, Lifesciences). All the results for fluorescence-based estimation of caspase-3 activity and the fold change relative expression of *HzCaspase-3* and *HzCaspase-8* were graphed and analyzed by unpaired t-test for individual comparisons between the treated and untreated cells, using the software GraphPad Prism 8.0.

### 4.8 Gene reporter assays

#### Luciferase assay in Wild type and Knock-out HzAW1 cells

To verify that OATP74D of *H. zea* acts as an ecdysone importer, an *in vitro* approach based on luciferase was employed. Specifically, the HzAW1^WT^ and HzAW1^ΔOatp74D^ cell lines were transfected with 1μg of the plasmid ERE- b.act.luc [50] in 6-well plates using ESCORT IV transfection reagent, following the instructions of the manufacturer.

Both cell lines were incubated for 72 hrs after transfection, after which 100μl of the transfected cells were seeded into 48-well plates and incubated for 2-3 hrs, followed by treatment with 0.1μM and 1μM of 20-ΗΕ (TCI chemicals, #1480). Twenty-four hours post treatment the cells were lysed and analyzed for luminescence using the Luciferase Assay system (Promega; Cat #E1500). Normalization among different technical replicates and conditions was carried out by normalizing the relative luminescence units (RLUs) against total protein content (calculated with the Bradford protein assay, BioRad). Each condition was measured in quadruplicates and each experiment was performed at least twice.

#### Luciferase assay in HzAW1^ΔOatp74D^ stably overexpressing HaOATP74D and SfOATP74D

For the OATP74D overexpression experiments, piE1:zeocinBmAc3-empty, piE1:zeocinBmAc3-*HaOatp74D,* piE1:zeocinBmAc3-*SfOatp74D,* or piE1:zeocinBmAc3-*DmOatp74D* were transfected along with ERE-b.act.luc plasmid in the HzAW1^ΔOatp74D^ cell line. The *DmOatp74D* and empty vector were used as positive and negative control respectively. Three days later 200 μl of the transfected cells were seeded into new 6-well plates treated with 0.01% poly-L-Lysine (Sigma), followed by selection with 1mg/ml of Zeocin (Invitrogen). The medium was refreshed every 4 days while selective concentration was reduced to 500μg/ml after 4 weeks of selection. Furthermore, a similar procedure was followed for the piE1:zeocinBmAc3-*HaOatp74D-V5* or piE1:zeocinBmAc3-*SfOatp74D-V5*, which both bear a V5 epitope tag at the C-terminus of the protein, in order to validate the expression of OATP74D in HzAW1^ΔOatp74D^ cell line. Validation of *HaOatp74D* and *SfOatp74D* was performed with Western blot and immunofluorescence, as described below.

After propagating the cell lines of each genotype, the cells were tested for responsiveness to 20-HE with the luciferase assay, following the same procedure as previously described. Approximately 10^5^ cells were seeded in 48- well plates coated with 0.01% poly-L-lysine (Sigma), followed by treatment with 20-HE overnight and were then tested for luciferase expression. Each condition was measured in eight independent technical replicates and each experiment was performed at least twice.

### 4.9 Western Blot and Immunofluorescence

For western blots, cell lines stably over-expressing *HaOatp74D-V5* and *Sf-Oatp74D-V5* were harvested and lysed with RIPA lysis buffer (50mM Tris-HCl pH 8.0, 150mM NaCl, 0.5% Sodium-Deoxycolate, 0.1% SDS and 1% NP-40) supplemented with 1X cocktail Protease Inhibitors (Sigma-Aldrich) and 1mM PMSF, followed by centrifugation for 10min at 4°C at 6,000g. Protein concentration was measured with Bradford assay (BioRad). Approximately 30μg of total protein was loaded onto 10% SDS-PAGE and subsequently transferred to a nitrocellulose membrane. A mouse anti-V5 antibody (Cell signaling) was used at a dilution of 1:2500 in 1% milk dissolved in 1X TBST buffer for detection of either HaOATP74D-V5 or SfOATP74D-V5 proteins. Anti-beta tubulin (Santa-Cruz) was also used at a 1:1000 dilution as loading control.

Cells over-expressing the epitope tagged lepidopteran OATP74D were used for immunostaining. Specifically cells were incubated on round shaped coverslips in 24-well plates. The cells were washed with 1X PBS and blocked for 1hr at room temperature with PBT solution, containing 2% BSA and 0.1% Triton-X100 in 1X PBS. The cells were incubated with 1:250 of primary antibody (mouse anti-V5) diluted in the blocking solution for overnight at 4°C. The cells were incubated with 1:1000 of anti-mouse secondary antibody conjugated with Alexa Fluor 555 for 1hr at room temperature. Nuclei were counterstained with DAPI and mounted with Vectashield Antifade mounting medium. Samples were observed using a Leica SP8 Inverted confocal microscope.

### 4.10 Cell based screening assay for inhibitors

To analyze the potency of broad-spectrum inhibitors of organic anion transporters to inhibit the function of lepidopteran OATP74D, HzAW1^ΔOATP74D^ cells stably overexpressing the lepidoptera OATP74D were pre-treated with several concentrations of telmisartan (Sigma-Aldrich) or rifampicin (Sigma-Aldrich) in the presence of 0.1μM 20-HE. Both inhibitors were tested at concentrations that do not impact cellular viability using the luciferase assay described above. Results were analyzed using the one-way ANOVA statistical test with Dunnet’s multiple comparison test.

## Acknowledgements

The authors would like to thank Dr. Cynthia L. Goodman (Biological Control of Insects Research, U.S, Department of Agriculture, Agriculture Research Service) for providing the RP-Hz-GUT-AW1 cell line. Moreover. the authors thank Ioannis Livadaras for performing the injections in *S. frugiperda* eggs.

---

**S1 Fig.**
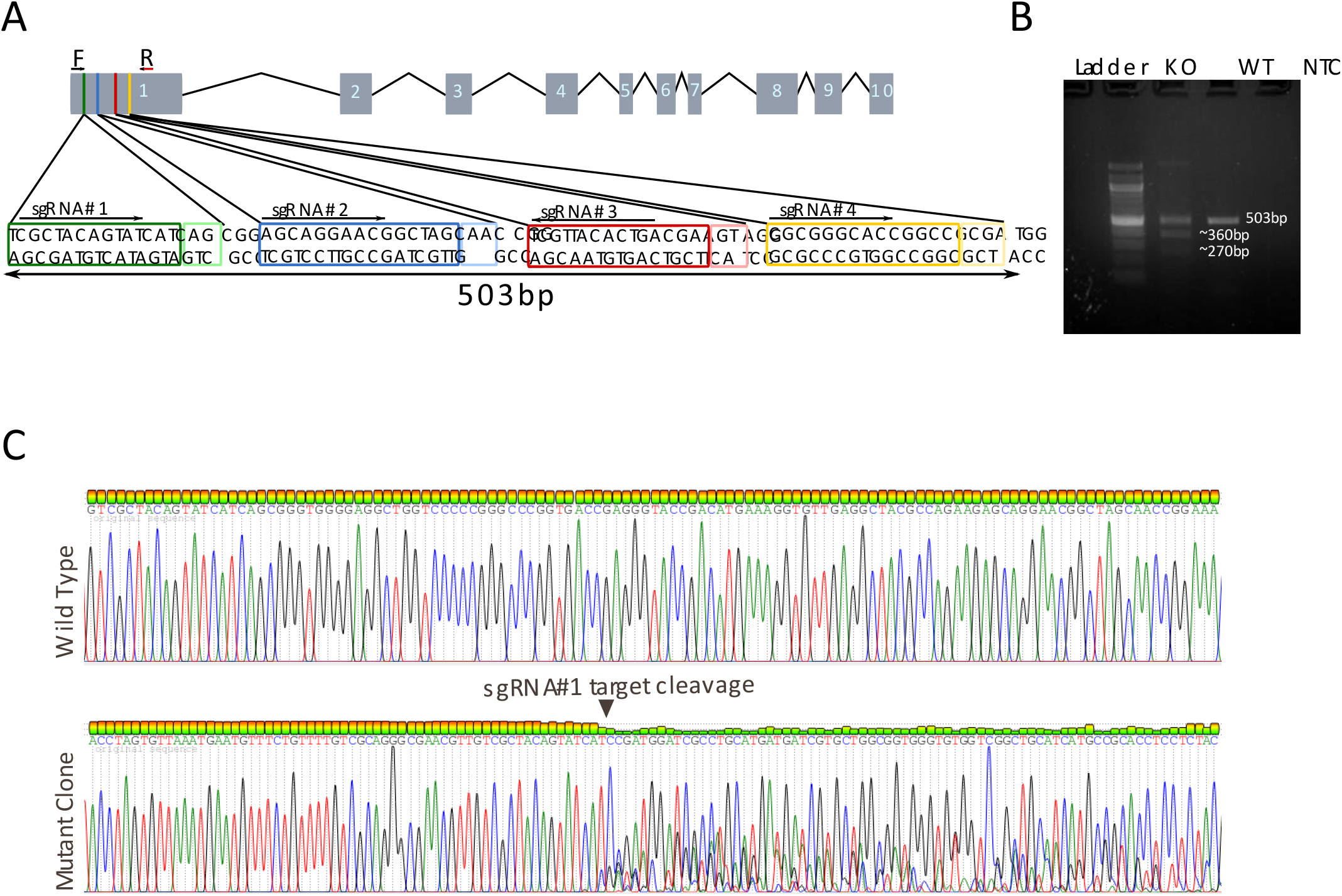
In vivo Characterization of Oatp74D in S. frugiperda. (A) Schematic representation of the SfOatp74D gene consisting of 10 exons. Four sgRNAs (#1, #2, #3 and #4)were designed to target the first exon of the gene spanning a region of 503bp. F and R indicate the forward and reverse primers respectively used in PCR for diagnostic reasons. (B) Diagnostic PCR screening yielding three fragments corresponding to a wild type (503bp product) and two deletions (360bp and 270bp) in CRISPR injected eggs; WT and KO indicate wild type and knock-out, NTC: non- template control. C) Sequencing chromatogram of region proximal to sgRNA#1 in wild type and mutant clone, indicating the disruption of the chromatogram downstream of the target region.

**S2 Fig.**
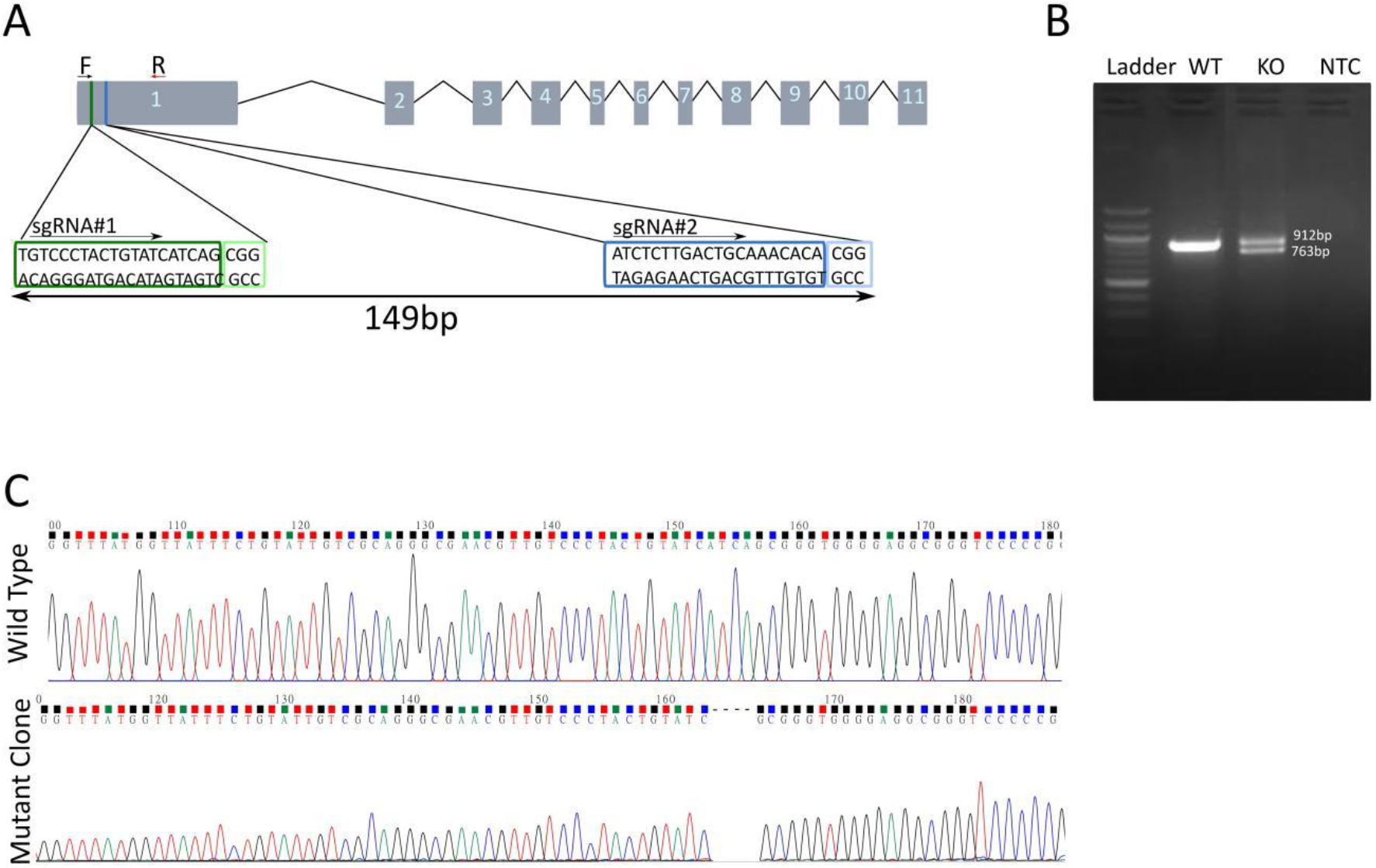
Characterization of Oatp74D in HzAW1 cell line. (A) Schematic representation of the HzOatp74D gene consisting of 11 exons.Two sgRNAs (#1 and #2) designed to target the first exon of the gene spanning a region of 149bp. F and R indicate the forward and reverse primers respectively used in PCR for diagnostic reasons. (B) Diagnostic PCR indicating the expected deletion of 149 bp after transfection of HzAW1 cells; WT and KO indicate the wild type and knock-out cells, NTC: non-template control. (C) Sequencing chromatogram of region proximal to sgRNA 1 in wild type and mutant clone, indicating the deletion of the 4 bp (5’-ATCA-3’).

**S3 Fig.**
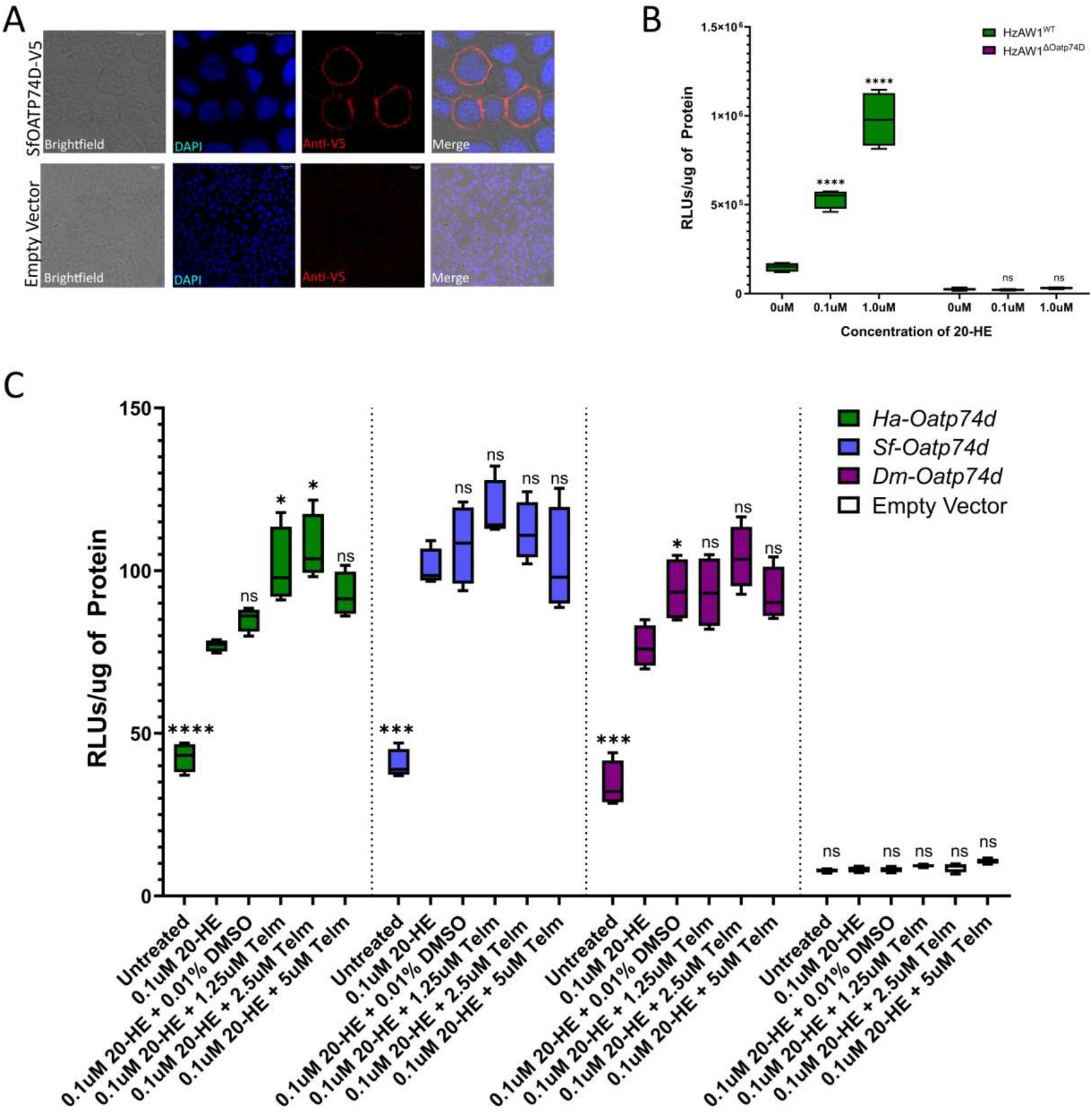
(A) Subcellular localization of SfOATP74D and HaOATP74D tagged with V5 epitope in transiently transfected Sf-9 cells. Blue indicates DAPI that counterstains nuclei, while Red indicates anti-V5, scale bar 20μM. (B) Luciferase assay in HzAW1WT and HzAW1ΔOATP74D cell lines after treatment with 0.1μΜ and 1μΜ 20-HE post transfection with the ecdysone responsive luciferase contruct. Asterisks indicate statistically significant differences between the treated and the untreated groups calculated with one-way ANOVA and post-Dunnet’s test, ****p<0.0001. (C) Inhibition analysis of lepidoptera OATP74D by Telmisartan in stable cell lines. The cell lines were incubated with 0.1uM of 20-HE in the presence of serial concentrations of Rifampicin (1.25μΜ, 2.5μΜ and 5μΜ) or DMSO (negative control). Each group is represented by eight different technical replicates. Asterisks indicate statistical significance between the different conditions versus treated only with 20-HE calculated with one-way ANOVA followed by post- hoc Dunnett test, *p<0.048, ***p<0.0008. ns: non-significant

**S4 Fig.**
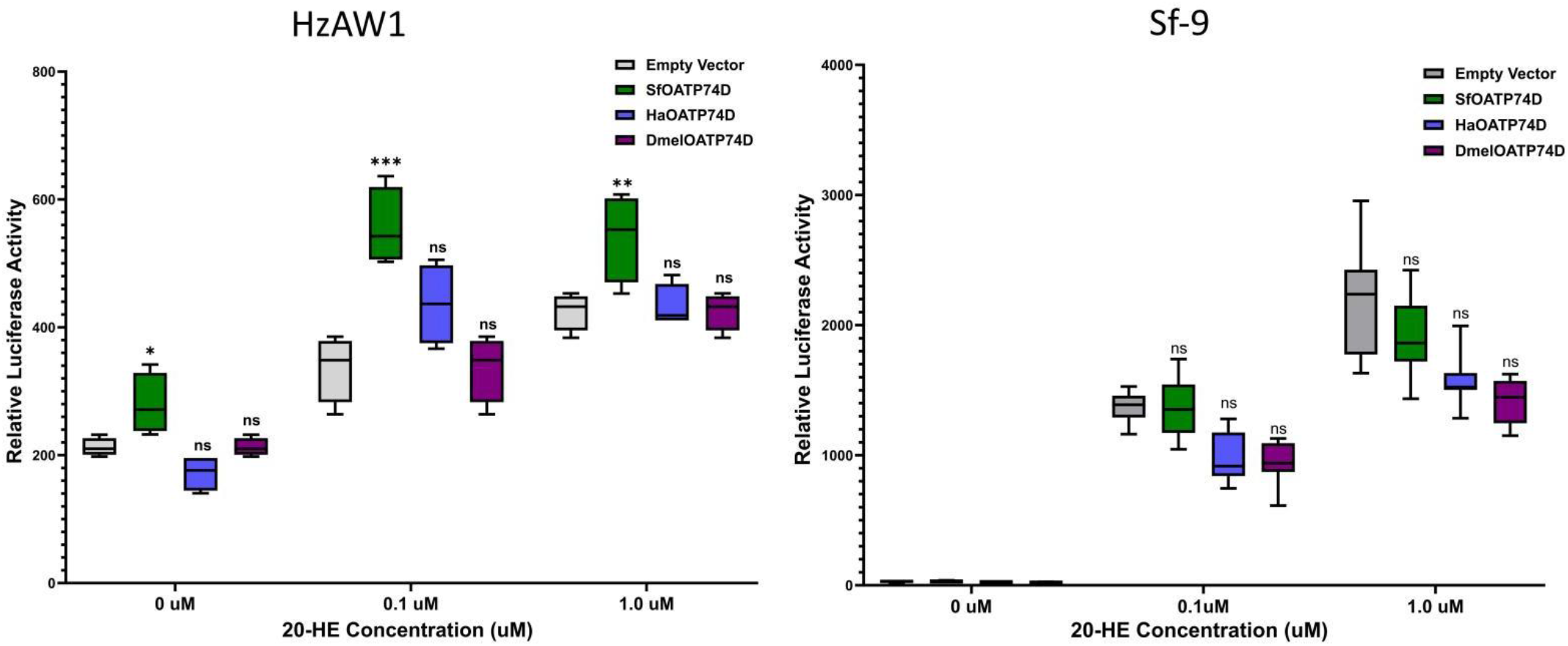
Functional characterization of lepidopteran OATP74D transiensly transfected in HzAW1WT (left) and Sf-9 cells (right). Cells were treated with several concentrations of 20-HE 72hrs post transfection and tested for luciferase expression 24hrs post treatment. Each group consists of four technical replicates and statistical significance was calculated by one-way ANOVA with post-Dunett’s test comparing cells overexpressing OATP74D with cells transfected with empty vector, *p<0.033, **p<0.0012, ***p<0.0008.

**S1 Table.**
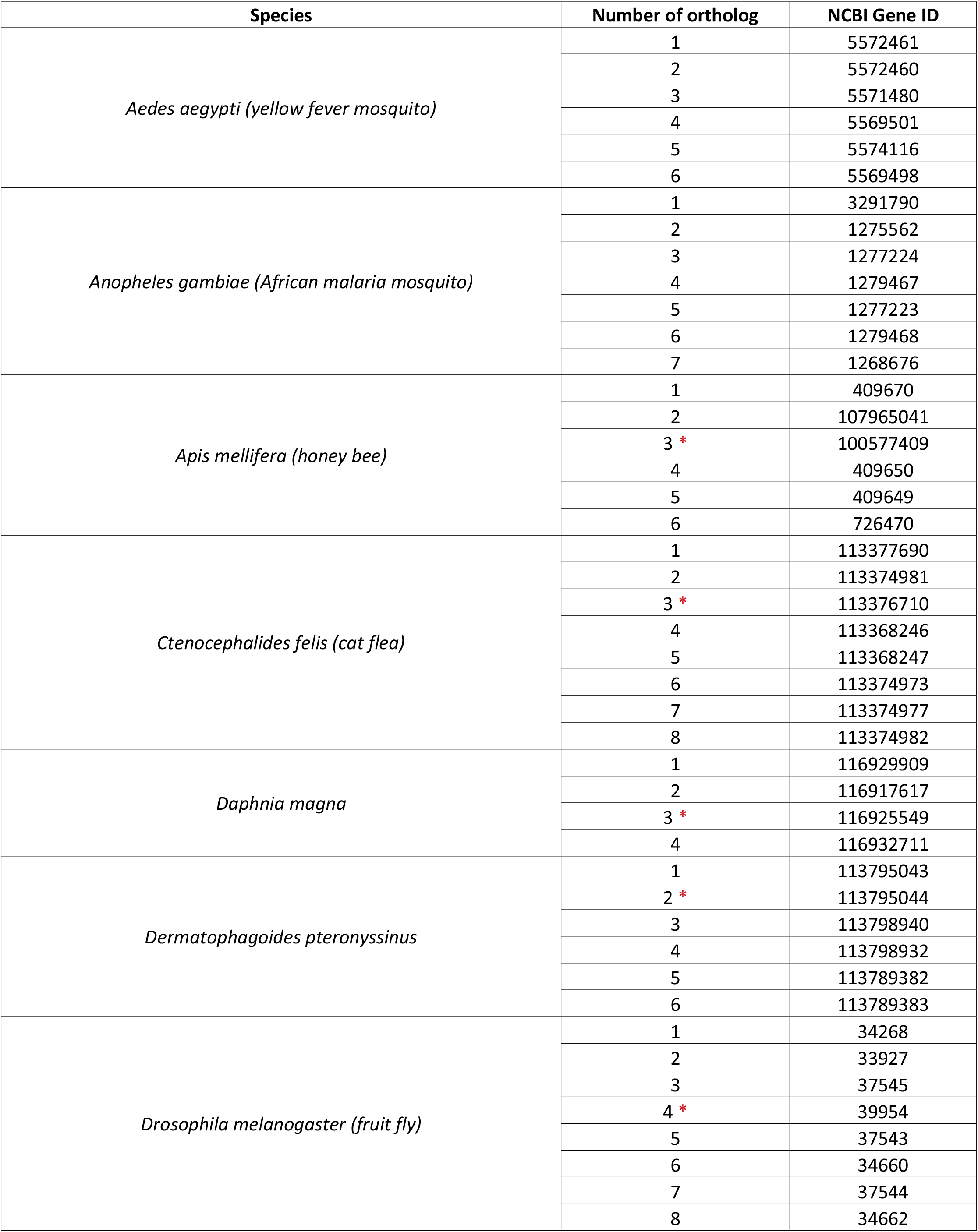
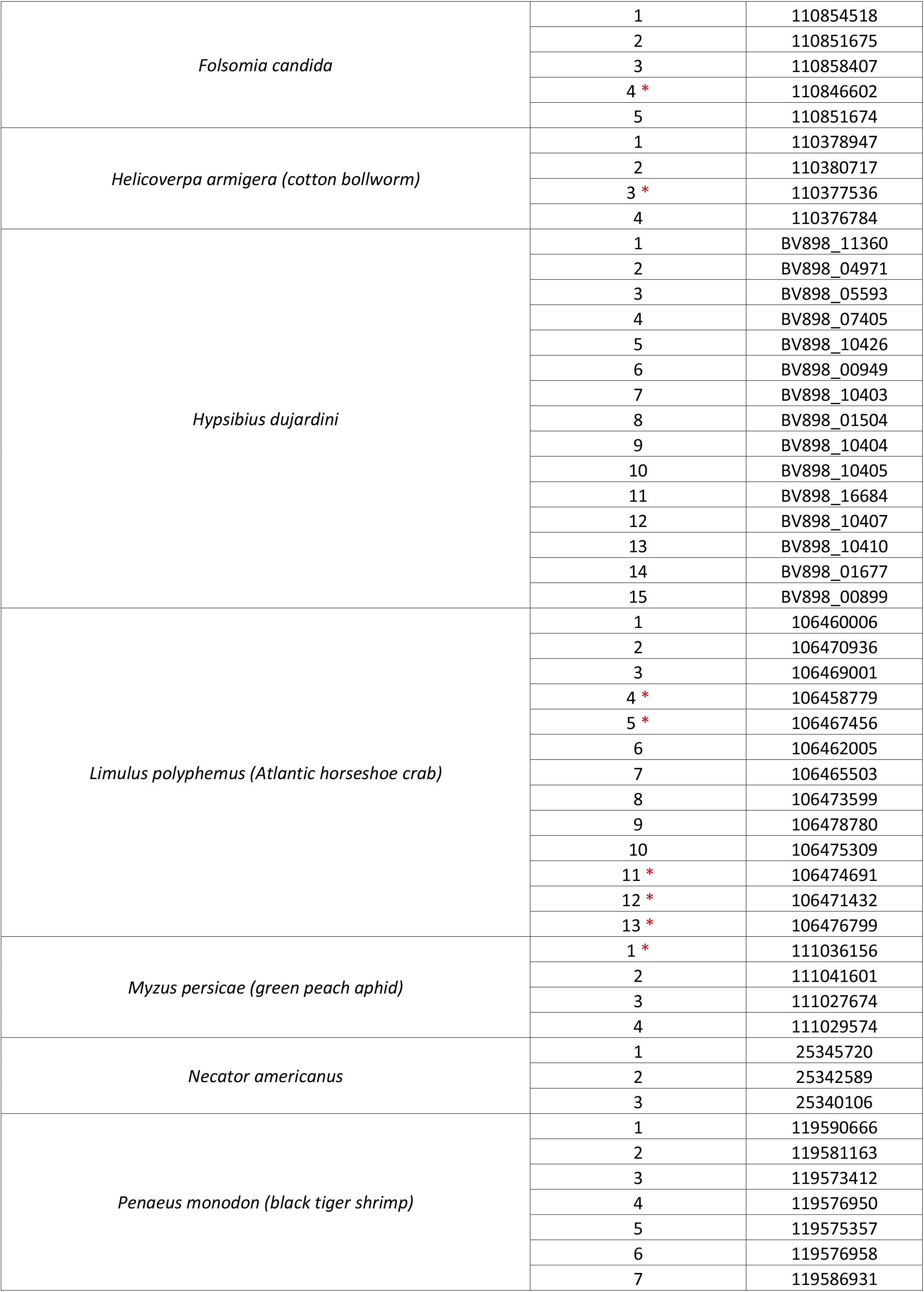
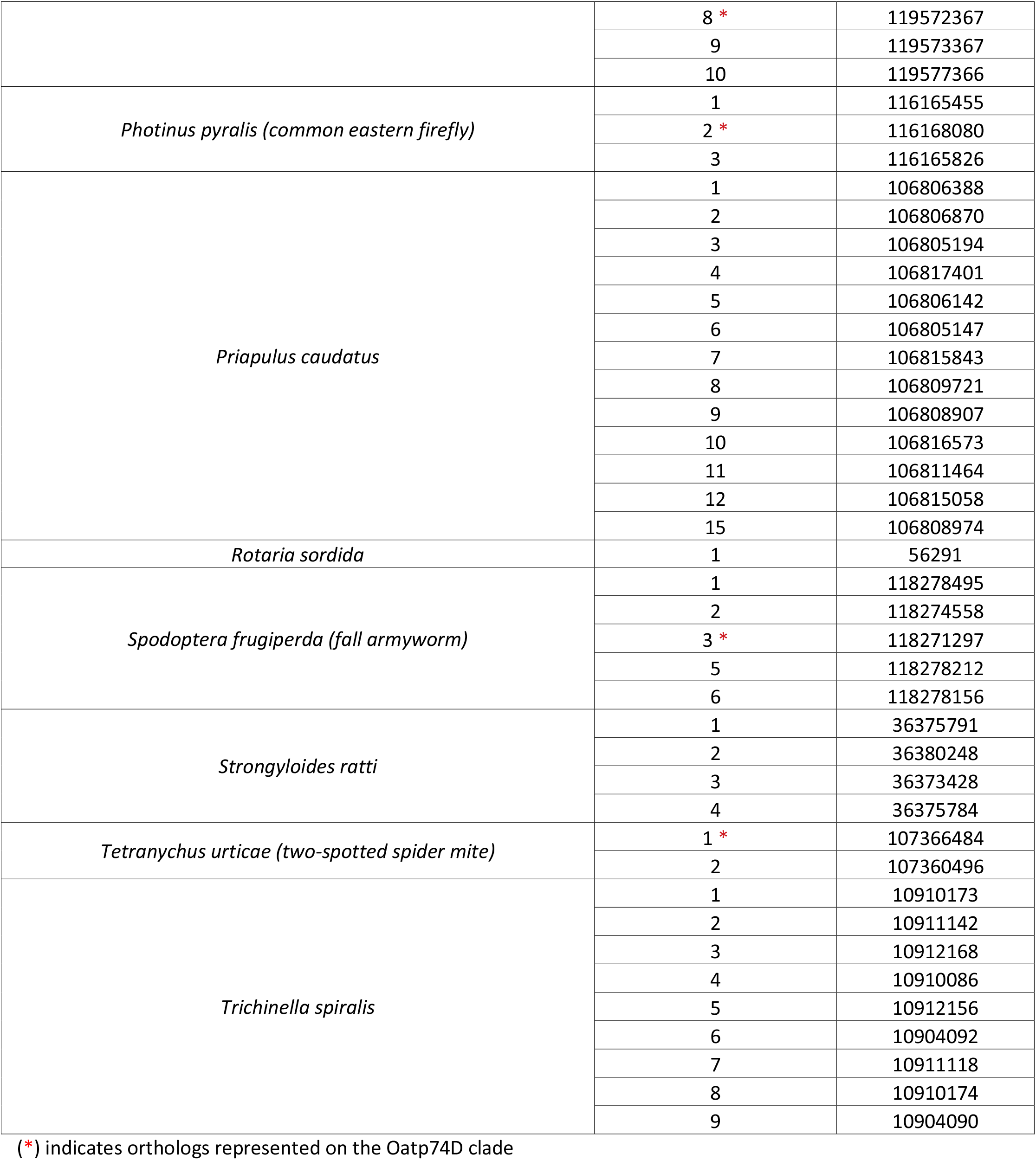
List of GenBank accession numbers of the proteins used for the construction of the phylogenetic tree.

**S2 Table.**
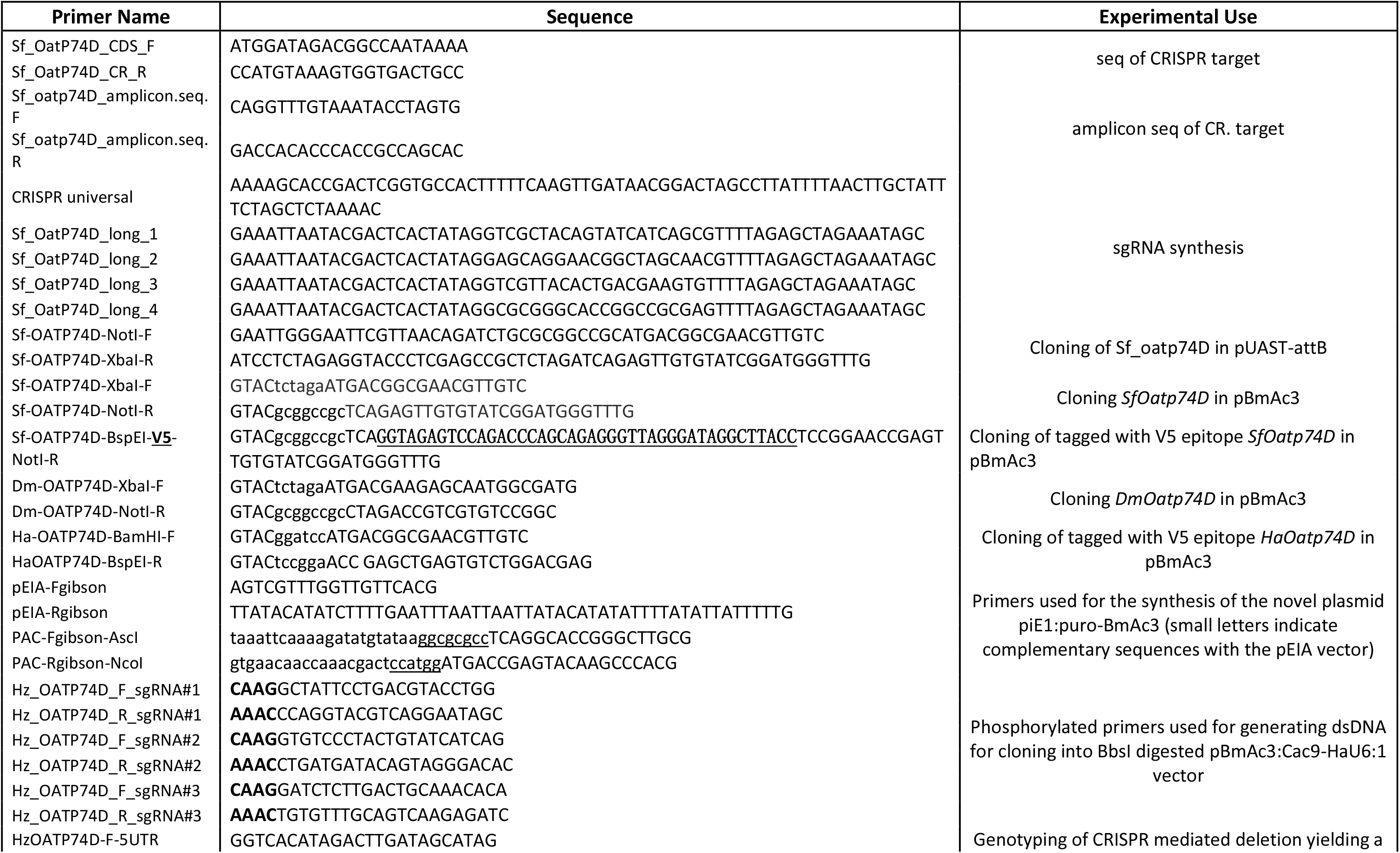
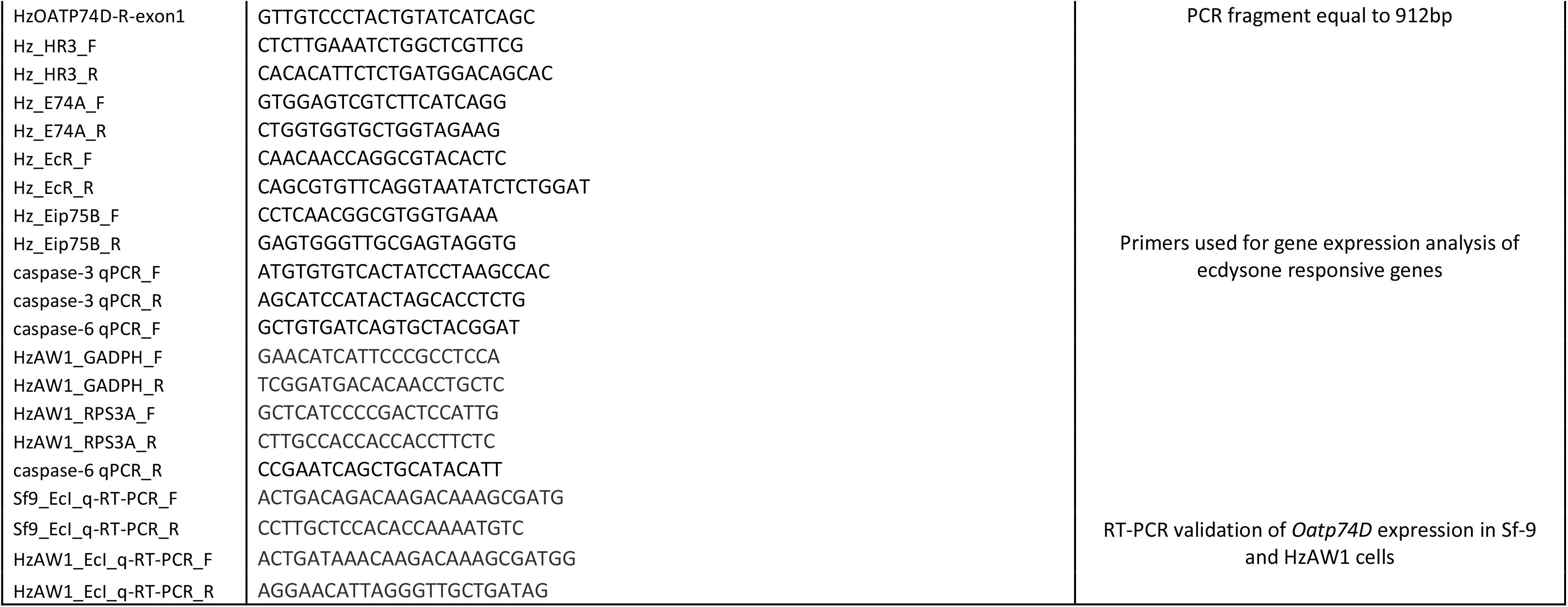
List of primers used in this study.

**S3 Table.**
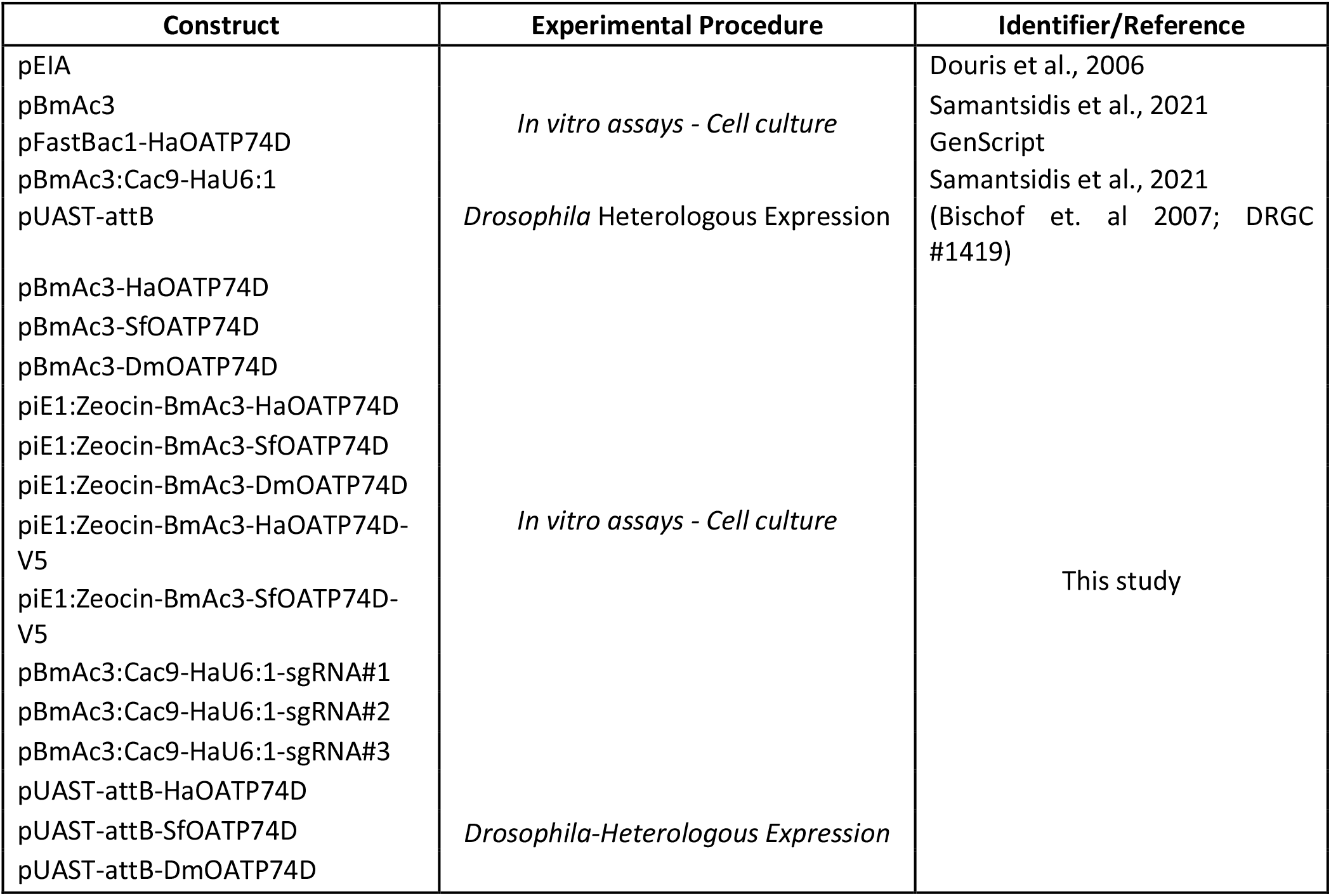
Table. List of plasmids used in this study.

**S4 Table.** Mortality scores monitored post injection of *S. frugiperda* targeting the *Oatp74D* or *Scarlet* genes.

| Genotype | Plate | Eggs | Hatched | Hatching<br>Rate<br>Percentile | L1 | L5 | L1-L5<br>Percentile | Overall<br>Mortality | Survival% | Average<br>Hatching<br>Rate | Average L1-L5 Trans. | Average<br>Surviv. |
| --- | --- | --- | --- | --- | --- | --- | --- | --- | --- | --- | --- | --- |
| *Oatp74D* | 1 | 87 | 10 | 11.49425287 | 10 | 0 | 0 | 0 | 0 | 10.2433881 | 25.09920635 | 25.09920635 |
| *Oatp74D* | 2 | 86 | 16 | 18.60465116 | 16 | 2 | 12.5 | 2.325581395 | 0.125 |  |  |  |
| *Oatp74D* | 3 | 91 | 6 | 6.593406593 | 6 | 1 | 16.66666667 | 1.098901099 | 0.166666667 |  |  |  |
| *Oatp74D* | 1 | 83 | 7 | 8.43373494 | 7 | 3 | 42.85714286 | 3.614457831 | 0.428571429 |  |  |  |
| *Oatp74D* | 2 | 81 | 7 | 8.641975309 | 7 | 2 | 28.57142857 | 2.469135802 | 0.285714286 |  |  |  |
| *Oatp74D* | 3 | 78 | 6 | 7.692307692 | 6 | 3 | 50 | 3.846153846 | 0.5 |  |  |  |
| control | 1 | 62 | 38 | 61.29032258 | 38 | 33 | 86.84210526 | 53.22580645 | 0.868421053 | 38.14563663 | 85.60785868 | 85.60785868 |
| control | 2 | 67 | 36 | 53.73134328 | 36 | 34 | 94.44444444 | 50.74626866 | 0.944444444 |  |  |  |
| control | 3 | 77 | 26 | 33.76623377 | 26 | 24 | 92.30769231 | 31.16883117 | 0.923076923 |  |  |  |
| control | 1 | 80 | 25 | 31.25 | 25 | 20 | 80 | 25 | 0.8 |  |  |  |
| control | 2 | 82 | 27 | 32.92682927 | 27 | 22 | 81.48148148 | 26.82926829 | 0.814814815 |  |  |  |
| control | 3 | 88 | 14 | 15.90909091 | 14 | 11 | 78.57142857 | 12.5 | 0.785714286 |  |  |  |

**S5 Table.** P-values of Student’s t-test for un-paired comparisons in gene expression analysis between the HzAW1WT and HzAW1*ΔOatp74D* cells.

|  | *HzEcR* | *HzHr3* | *HzEip74A* | *HzEip75B* |
| --- | --- | --- | --- | --- |
| **6hrs** | 0.0053 | 0.0032 | 0.0046 | 0.4597 |
| **9hrs** | 0.0048 | <0.0001 | 0.0321 | 0.0011 |
| **12hrs** | 0.0031 | 0.0002 | 0.0451 | 0.003 |
| **24hrs** | 0.0379 | <0.0001 | 0.026 | 0.0744 |

